# Single-Cell Spatial Proteomics Uncovers Molecular Interconnectivity among Hallmarks of Aging

**DOI:** 10.64898/2026.02.26.708335

**Authors:** Seungmin Yoo, Christopher Young, Liying Li, Lingraj Vannur, Junming Zhuang, Fan Zheng, Meiying Wu, Julie K. Andersen, Chuankai Zhou

**Author notes:** Equal contribution (Seungmin Yoo, Christopher Young, Liying Li, Lingraj Vannur).

## Abstract

Aging is accompanied by conserved hallmarks including genomic instability, epigenetic alterations, loss of proteostasis, and mitochondrial dysfunction, but how these processes emerge and become mechanistically linked remains unclear. Here we leverage a proteome-wide, single-cell, subcellular atlas of protein expression, localization, and aggregation across yeast replicative aging to map hallmark-linked remodeling in its spatial context. We identify hundreds of previously unappreciated molecular changes that underlie major hallmarks of aging and show that hallmark phenotypes frequently manifest as compartment-specific erosion of spatial confinement, relocalization, and aggregation. 91.6% human orthologs of these hallmark-linked yeast proteins also change during human aging. Integrating these spatial phenotypes reveals many molecular connections linking different hallmarks. Temporal analysis suggests that disorganization of nucleolar ribosome biogenesis, proteostasis decline, and mitochondrial dysfunction precede other hallmarks. Together, our findings substantially deepen the molecular underpinnings of aging hallmarks and provide a framework for linking them into a hierarchical sequence of cellular failures.

## Introduction

Aging is characterized by a progressive loss of molecular organization and physiological resilience^1,2^. The hallmarks of aging—including genomic instability, epigenetic alterations, loss of proteostasis, and mitochondrial dysfunction—have provided a powerful conceptual framework for organizing these changes into conserved molecular categories^1,2^. However, most studies have examined these hallmarks in isolation, focusing on individual pathways or molecular classes. As a result, we still lack an integrated understanding of how distinct hallmarks arise, how they interact, and which molecular events bridge them. A comprehensive map linking these hallmarks has remained elusive.

Progress toward this goal has been limited by the absence of datasets that capture the aging proteome in its full spatial and quantitative context. Most molecular studies focus on transcriptomic or bulk-proteomic averages, which obscure subcellular organization and cell-to-cell variability^3,4^. Conversely, prior imaging studies provided valuable spatial context but were largely limited to individual proteins or specific organelles^5–8^, which constrained their ability to capture proteome-wide remodeling and to place pathway-specific changes within a global framework. Because many cellular defects underlying the hallmarks of aging—such as nucleolar enlargement, organelle morphological alterations, loss of compartmentalization, protein re-localization, and aggregation—are inherently spatial, their emergence and propagation can only be captured through proteome-scale imaging capable of resolving localization, morphology, and abundance simultaneously in single cells. The absence of such a comprehensive microscopy dataset represents a major gap in aging biology, preventing a spatially resolved understanding of how the different hallmarks of aging emerge during aging.

In the companion study (Yoo et al., 2026a), we addressed this gap by constructing a proteome-wide, single-cell atlas of protein expression, subcellular organization, and aggregation behavior during yeast replicative aging, using a library of strains with proteins endogenously and seamlessly tagged with mNeonGreen (mNG). That study established the experimental and computational platform for quantitative three-dimensional imaging and analysis of more than 94% of the yeast proteome and >90 million cells across replicative aging, captured with high-content confocal microscopy. Using a deep-learning–based analysis pipeline, we quantified concentration, localization, and aggregation states of proteins at single-cell and single-age resolution, generating an unprecedented dataset for exploring proteome dynamics. The resulting dataset captured widespread protein localization shift, reduction of protein local concentration, loss of organelle scaling relationships, and remodeling of protein interaction network, while revealing intrinsic structural determinants underlying the fate of proteins during aging. Collectively, these efforts generated a comprehensive reference resource for exploring the spatial and temporal organization of the aging proteome.

In this study, we leverage this single-cell spatial atlas to dissect the molecular changes that underlie—and connect—conserved hallmarks of aging. By integrating spatiotemporally and subcellularly resolved measurements across the proteome, we uncover hundreds of previously unappreciated age-associated remodeling events across ribosome biogenesis, chromatin regulation, proteostasis, RNA metabolism/translation, nutrient transport, and mitochondrial metabolism, thereby refining the molecular definition of multiple hallmarks. These compartment-resolved phenotypes reveal a dense network of molecular connections among hallmark-associated processes. Temporal trajectories indicate a progressive deepening of dysfunction from early ages, with nucleolar disruption, proteostasis decline, and mitochondrial dysfunction accelerating ahead of other hallmark programs. Together, our data support a view of aging as an interconnected proteome reorganization rather than independent failures, and highlight the value of subcellular spatiotemporal proteomics for revealing links between hallmarks that are difficult to infer from RNA-seq or bulk mass spectrometry.

## Results

### Age-associated molecular changes underlying genomic instability in the nucleus and mitochondria

#### Nuclear DNA

Genome integrity and stability are increasingly challenged with age across all species^2^. Consistent with this, we observed an age-associated increase in single-stranded DNA (ssDNA) foci in the nucleus as visualized by RPA complex proteins (Figure 1A). Similarly, components of multiple DNA repair pathways—including homologous recombination (Rad52 and Rdh54), nucleotide excision repair (Rad1 and Cdc9), and base excision repair (Ogg1 and Apn1)—were significantly upregulated in a substantial fraction of the aged cells, consistent with a cellular response to accumulating DNA damage (Figure 1B). These DNA damages may suppress DNA replication, as MCM complex was markedly reduced in aged cells (Figure S1A). In addition to DNA damage, we observed significant loss of the spindle pole body (SPB) component Spc98, which could contribute to genomic instability (Figure 1C)^9^. Moreover, we observed frequent age-associated nuclear-shape abnormalities (∼30% of aged cells; Figure 1D) together with pronounced nuclear enlargement (Figure 1E), phenotypes reminiscent of the misshapen nuclei reported in mammalian aging/cellular senescence and in Hutchinson–Gilford progeria syndrome^10,11^.

**Figure 1.**
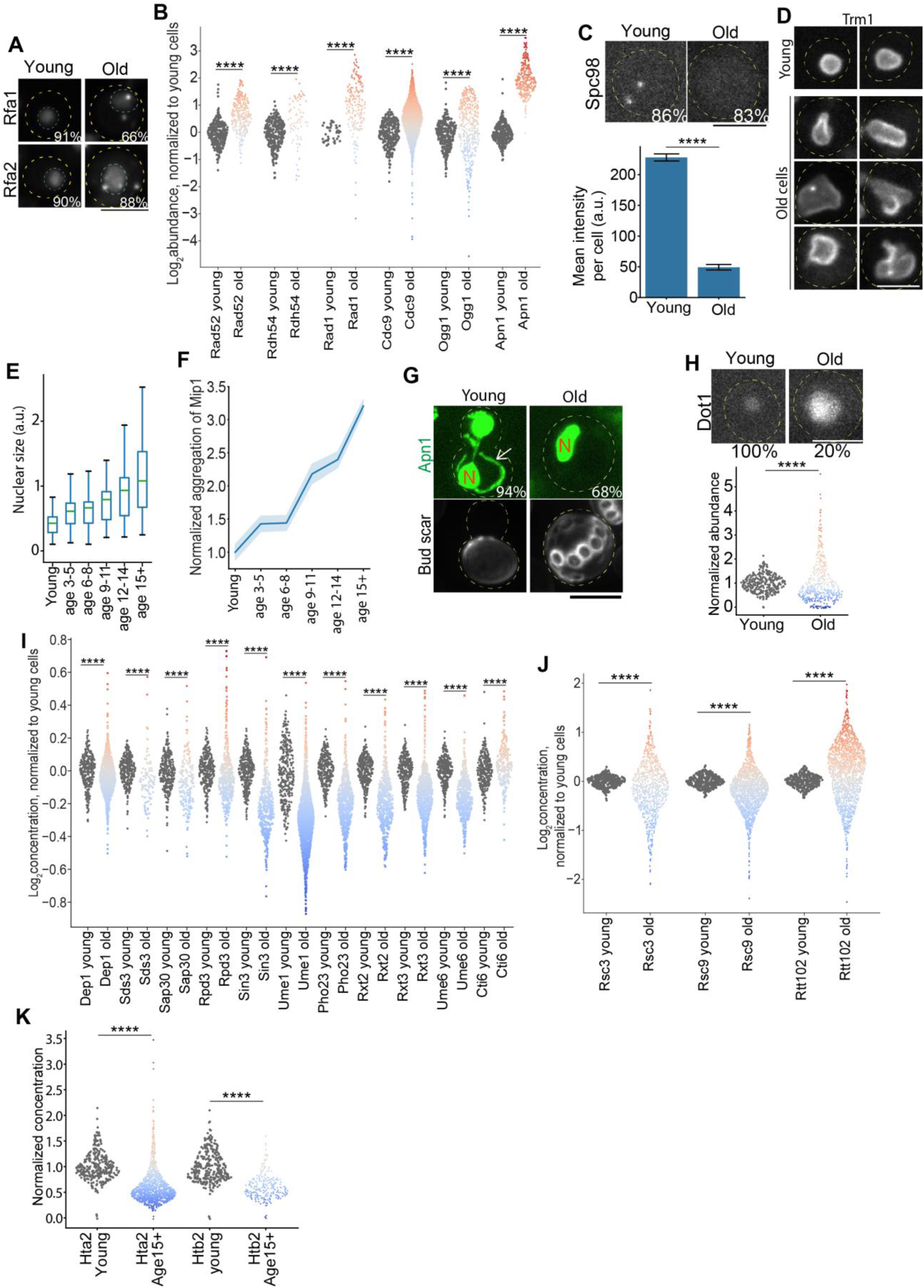
Age-associated molecular changes underlying genomic instability and epigenetic alterations. (**A**) Representative images of Rfa1/2 localization and foci during aging. The inset percentage indicates, for each age group, the fraction of cells exhibiting the phenotype shown in the representative images (same definition used in other figures). The cyan dashed line outlines the nucleus. (**B**) Quantification of protein abundance for DNA repair-related proteins in old cells (≥15 generations; same definition used in other figures). Each dot represents an individual old cell, with values normalized to the mean value of the corresponding protein in young cells. Dot color indicates fold change relative to the young-cell mean (red: increase; blue: decrease; same in other figures). (**C**) Representative images and quantification of Spc98 during aging. Shown are mean and standard error. (**D**) Representative images showing nuclear-envelope morphology across aging stages. Trm1, a nuclear-envelope–associated protein, is shown to visualize nuclear-envelope structure. (**E**) Quantification of nuclear size across different aging stages. Dig2, a nuclear protein, was used for this quantification. Shown are mean and standard deviation. (**F**) Quantification of protein aggregation levels of Mip1 across different aging stages. Shown are mean and standard error (shade). (**G**) Example cells and quantifications of Apn1localization at different aging stages. Arrow point to mitochondrial signal. N, nucleus. (**H**) Example cells and quantifications of Dot1 at different aging stages. (**I-K**) Quantification of protein concentration for Rpd3L complex (I), RSC complex (J), and histone proteins (K) in old cells. Scale bar: 5μm. See table S5 for number of cells quantified.

In addition to general genomic instability, another highly conserved aging phenotype is nucleolar morphological changes and rDNA instability^12–16^. We observed similar nucleolar remodeling during replicative aging, accompanied by age-dependent aggregation of select nucleolar proteins (Figure S1B, see also “loss of proteostasis” and “ribosome biogenesis defect” below). Beyond these morphological changes, replicative aging in yeast is linked to increased ribosomal DNA (rDNA) instability and the accumulation of extrachromosomal rDNA circles (ERCs)^17,18^. This has been attributed to age-associated declines in NAD⁺ and reduced Sir2 activity, which compromise rDNA silencing and promote recombination at the rDNA locus, thereby driving ERC formation^19–22^. Consistently, we observed reduced abundance of Nma2, a key NAD⁺ biosynthetic enzyme (Figure S1C). A recent work suggests that only a subset of aged cells exhibits rDNA silencing defects^23^. Consistent with this cell-to-cell heterogeneity, we observed an age-dependent reduction of Sir2 in a fraction of old cells, whereas other aged cells showed increased Sir2 abundance (Figure S1D). We also found that Fob1—central to replication fork arrest, rDNA recombination, and ERC generation^24^—became delocalized from the nucleolus in ∼30% of old cells, displaying reduced abundance and/or a smeared nuclear distribution relative to young cells (Figure S1E). Sir2 may be further perturbed in old cells that lost Ulp1 (Figure S1F), a major SUMO protease, given that SUMOylation modulates Sir2 localization and transcriptional silencing^25^. Together, these changes are consistent with weakened rDNA silencing and increased rDNA recombination in a subset of old cells during replicative aging^23^.

#### Mitochondrial DNA (mtDNA)

Because mtDNA encodes essential subunits of oxidative phosphorylation (OXPHOS), mitochondrial genome instability provides a mechanistic link between genomic instability and mitochondrial dysfunction—two hallmarks of aging. mtDNA instability involves changes in both mtDNA quantity (copy number) and quality (mutations/heteroplasmy)^2,26,27^. In line with reports of age-associated mtDNA loss^26,28^, we observed aggregation of the mitochondrial DNA polymerase γ, Mip1, in aged cells, suggesting compromised activity of this essential replication factor (Figure 1F). We also detected aggregation of Mhr1 (Figure S1G), a mitochondrial recombinase implicated in recombination-dependent/rolling-circle replication, mtDNA segregation, and repair of mitochondrial double-strand breaks^29–31^. Together, these defects could contribute to the progressive loss of mtDNA with age^26,28^.

mtDNA instability also arises from DNA damage caused by oxidation and other chemical modifications generated by the unique metabolic environment within mitochondria^32–35^. Base excision repair (BER) is a principal pathway for resolving small base lesions arising from oxidation, alkylation, deamination, and methylation^36^. Notably, several BER factors—including DNA glycosylases (Ogg1, Ung1, Ntg1, Ntg2), the 3′-repair diesterase Apn1, and DNA ligase Cdc9—localize and function in both the nucleus and mitochondria; in young cells, a small portion of each protein is imported into mitochondria to maintain mtDNA integrity^37–42^. Disrupting mitochondrial import of these BER enzymes is known to increase mtDNA mutagenesis and accelerate mtDNA depletion^37,43^. Strikingly, we found that aged cells selectively lost these BER enzymes from mitochondria (Figure 1G, S1H–M), suggesting age-dependent defects in mitochondrial import and/or mitochondrial proteostasis. Because several of these enzymes increased with age (Figure 1B) and only a small fraction was mitochondrially localized, this organelle-specific depletion would be missed by bulk measurements that average across compartments and cells. Together, this compartment-specific loss of mitochondrial BER enzyme import/retention provides a mechanism by which age-associated proteostasis and protein-import defects may contribute to mtDNA instability and mitochondrial dysfunction (see also “loss of proteostasis” and “mitochondrial dysfunction” below).

### Epigenetic alterations: remodeling of histone and chromatin modifiers

Replicative aging in yeast is accompanied by extensive remodeling of chromatin structure and histone modifications, affecting both transcriptional fidelity and genome stability^44–46^. Accordingly, we mined the atlas for molecular changes underlying these epigenetic alterations, including regulators of rDNA/telomeric silencing, HDAC complexes, histone abundance, and ATP-dependent chromatin remodelers.

Dot1 is the sole H3K79 methyltransferase, and hypermethylation of H3K79 has been reported in old yeast and in aging human neurons^47,48^. Dot1-dependent H3K79 methylation promotes DNA damage responses and antagonizes Sir protein association with chromatin, thereby reducing SIR activity at rDNA repeats and telomeres^49–51^. We observed an increased frequency of Dot1-high cells among old cells: although the population-average abundance of Dot1 declined, a subset of aged cells exhibited elevated Dot1 levels (Figure 1H), a pattern consistent with age-associated H3K79 hypermethylation^47,48^ and weakened Sir-dependent rDNA/telomeric silencing^49–51^. In parallel, we observed a reduction in the Cohibin components Csm1 and Lrs4 (Figure S2A), which tether rDNA to the nuclear periphery to support rDNA silencing and suppress recombination^52,53^. We observed a similar age-associated loss of Sir3, a core SIR complex subunit required for heterochromatin formation and transcriptional silencing at the HML/HMR mating-type loci and telomeres (Figure S2A)^54–57^, suggesting that telomere-proximal repression may be compromised in aged cells. Consistent with this prediction, analysis of published transcriptomic data^58^ indicates that many subtelomeric genes were transcriptionally upregulated (Figure S2B). Together, these molecular changes suggest a route by which epigenetic alteration may promote genomic instability during aging.

Histone deacetylase (HDAC) complexes undergo subunit-specific remodeling with age. Yeast Class I HDACs include Rpd3 (in the Rpd3L and Rpd3S complexes) as well as Hos1 and Hos2^59,60^. Within the Rpd3L complex, which primarily represses transcription at promoters, several subunits including Ume1, Pho23, Rxt2, and Rxt3 were reduced in old cells, suggesting altered complex composition and activity (Figure 1I) ^38,39^. Within the Rpd3S complex, which deacetylates transcribed regions to suppress cryptic transcription, Rco1 and Eaf3 decreased (Figure S2C), consistent with weakened repression of cryptic unstable transcripts (CUTs) and aligning with the conserved age-associated increase of CUTs^61–64^. The Set3C complex, containing Hos2 and Sif2, also declined and became cell-to-cell heterogeneous (Figure S2D). Given Set3C’s role in repressing meiotic programs, this erosion of repression may contribute to ectopic activation of sporulation genes in mitotically cycling aged cells (Figure S2E) ^61,65^. It is important to note that most of these sporulation-related genes were not detectable at the protein level; Dmc1—the meiosis-specific recombinase required for double-strand break repair—was the major exception and increased in aged cells (Figure S2F). Together, these observations support a model in which aging increases transcriptional noise and reduces repression fidelity rather than triggering true meiotic entry. Finally, the Class II HDAC Hda1 complex showed reduced and heterogeneous Hda1 levels in old cells (Figure S2G)^66^, which may further diversify transcriptional states within aged populations.

ATP-dependent chromatin remodelers are also affected by aging. We detected elevated levels of the SWI/SNF subunit Snf6, and an increase in multiple RSC components (e.g., Rsc3 and Rsc9) in some but not all aged cells (Figure 1J, S2H-I). These remodelers, along with shared components such as Rtt102, showed cell-to-cell variability in aged cells and could exacerbate transcriptional heterogeneity among aged cells by altering nucleosome positioning at promoters and nucleosome-depleted regions (Figure 1J)^44,66^. The reduction of histone concentration in the nucleus during aging likely further contributes to the epigenetic shift and increase in transcriptional noise (Figure 1K) ^44,61,66^. Interestingly, we observed increased abundance of Mtr4, an RNA surveillance helicase that promotes nuclear exosome–mediated RNA decay, as well as RNase III Rnt1, a dsRNA/hairpin-cleaving endonuclease, in some aged cells, potentially reflecting a compensatory response to increased antisense/aberrant transcription and dsRNA accumulation, consistent with reports in human senescence (Figure S2J-K)^67,68^. Together, these age-dependent alterations to histone modifiers and chromatin remodelers support a model in which multi-layered erosion of epigenetic regulation destabilizes transcriptional programs and genome integrity during aging.

### Age-associated molecular changes underlying autophagy impairment and loss of proteostasis

A primary hallmark of cellular aging is the progressive collapse of proteostasis, manifested as impaired protein translation, folding, degradation, and organelle/cargo turnover via autophagy^2,69^. In aged yeast cells, we observed striking disruptions in the cytoplasm-to-vacuole targeting (Cvt) pathway, marked by the accumulation of Ape1 and Ams1 puncta and persistent Atg19 and Atg11 foci (Figure 2A, S3A). These intermediates indicate inefficient delivery of Cvt cargo to the vacuole and are consistent with reported age-associated defects in vacuolar acidification and autophagy^5,70–72^. Their persistence in the cytosol highlights a bottleneck in vacuolar clearance capacity likely to exacerbate cytosolic proteotoxic stress. We also detected a more modest age-dependent increase in Atg20 and Atg21 puncta (Figure 2A), suggesting broader perturbation of trafficking modules that support autophagy.

**Figure 2.**
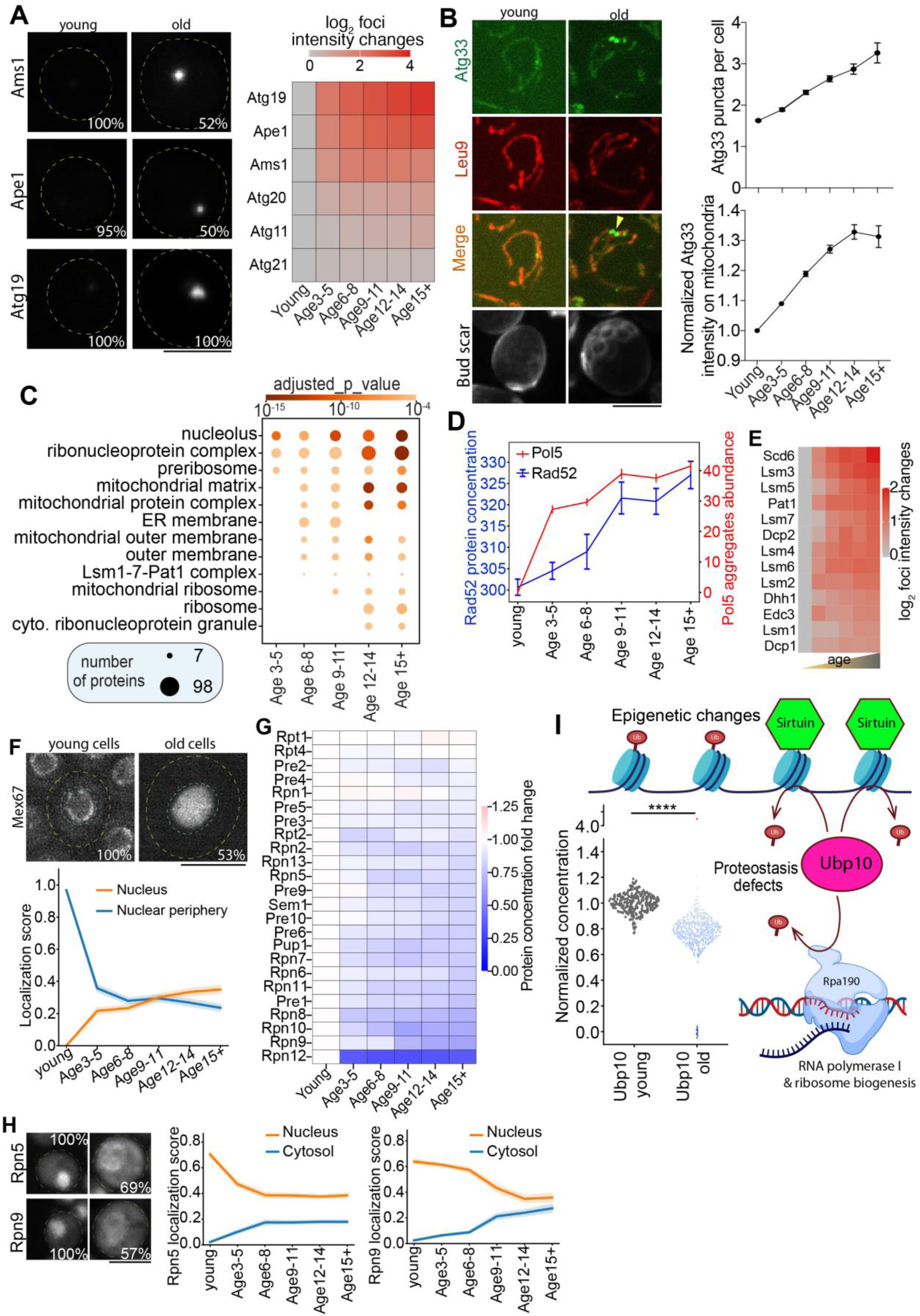
Age-associated molecular changes underlying disabled autophagy and loss of proteostasis. (**A**) Representative images and quantifications of puncta intensity changes for different proteins involved in cytosol-to-vacuole targeting (Cvt) pathway and autophagy during aging. (**B**) Representative images and quantifications for Atg33 in young and old cells with Leu9 as mitochondrial marker. White arrowhead indicates an example of Atg33 puncta on mitochondria. (**C**) GO term cellular compartment enrichment for the aggregating proteins during aging. (**D**) Quantification of age-associated Pol5 aggregation in the nucleus and increased Rad52 concentration (DNA damage response marker). Shown are mean and standard error. (**E**) Quantifications of puncta intensity changes for different P-body proteins during aging. (**F**) Representative images and quantifications of Mex67 localization changes during aging. Shown are mean and standard error (shade). The cyan dashed line outlines the nucleus. (**G**) Quantifications of protein concentration changes for different 19S subunits of proteasome during aging. (**H**) Representative images and quantifications of Rpn5 and Rpn9 localization changes during aging. Shown are mean and standard error (shade). (**I**) Schematic illustrating Ubp10’s dual functions, together with quantification of Ubp10 protein concentration in young and old cells (≥15 generations). Each dot represents an individual old cell, with values normalized to the mean Ubp10 level in the corresponding young cells. Dot color indicates fold change relative to the young-cell mean (red: increase; blue: decrease). Shown are mean and standard error (error bars or shade). Scale bar: 5μm. See table S5 for number of cells quantified.

Beyond these general defects in Cvt trafficking and core autophagy machinery, organelle-selective autophagy pathways were also compromised with age. Mitophagy selectively removes damaged mitochondria and requires Atg33, a factor for the selection or detection of damaged or aged mitochondria^73–75^. Notably, Atg33 accumulated and formed punctate foci on some mitochondria in aged cells (Figure 2B), a pattern suggestive of reduced or stalled mitophagy and accumulation of damaged mitochondria during aging^76^. Nucleophagy occurs at nucleus–vacuole junctions (NVJs), formed by the nuclear envelope protein Nvj1 and the vacuolar protein Vac8, where portions of the nucleus are engulfed by the vacuole^77^. Similarly, the NVJ tether protein Nvj1, but not Vac8, was diminished in most aged cells (Figure S3B), indicating loss of NVJs and thus reduced nucleophagy capacity.

In line with degradative defects of damaged proteins and organelles, our proteome-wide analysis revealed that protein aggregation becomes pervasive across multiple compartments during aging (Figure 2C, S3C-D, Table S1). Aggregation of numerous nucleolar proteins and mitochondrial chaperones/proteases—including Hsp78, Pim1, Cym1, and Oma1—highlights destabilization of organellar proteostasis networks (Figure 2C, S1B, S3E; see also “mitochondrial dysfunction” below). Pathway analysis of aggregated proteins further underscores broad susceptibility across processes including amino acid biosynthesis, mRNA decapping, rRNA processing, mitochondrial biogenesis, ATP synthesis, and ribonucleoside metabolism (Figure S3F). Importantly, compartments were not equally vulnerable: aggregation rose first in the nucleus and nucleolus, detectable as early as generations 3–5, preceding widespread aggregation in mitochondria, ER, and cytosol (Figure 2C). This early susceptibility positions the nucleolus and ribosome biogenesis machinery as a proteostasis weak point, with the potential to amplify downstream proteotoxic stress given the central role of ribosome production and translational control in maintaining proteome integrity^78^. Consistent with this idea, translational stress and impaired translational control are conserved features of aging, and interventions that improve translation fidelity can extend lifespan^79–81^. Notably, proteostasis disruption in the nucleus and nucleolus emerged early and was detected before changes in DNA damage response markers during aging (Figure 2D), suggesting that early nucle(ol)ar proteostasis failure is an upstream lesion that may contribute to downstream genomic instability.

Similar to proteostasis defects induced by environmental stresses, the age-associated collapse of proteostasis also induced cytosolic P-body accumulation, as evidenced by foci formation of RNA-binding proteins such as Scd6, Dcp1/2, and Dhh1 (Figure 2E), pointing to global translation repression in these aged cells^8^. P-body formation is consistent with translation repression that reduces protein folding load and shifts mRNAs from polysomes into dormant cytosolic mRNA granules when proteostasis is challenged^82^. This aligns with reports that aging is accompanied by translational arrest and increased ribosome pausing during elongation^79,80^. Consistently, we observed elevated Ssd1, a translational repressor (Figure S3G). Because Ssd1 activity has been linked to lifespan extension, its upregulation likely reflects a pro-longevity response to proteostasis challenge^83^.

In addition to cytosolic mRNA sequestration, abnormal nuclear RNA metabolism likely further contributes to translation and proteostasis defects. We observed redistribution of Mex67, a key mRNA export factor, from the nuclear pore complex (NPC) into the nucleoplasm (Figure 2F). Notably, this change coincided with similar relocalization of Nup2 (Figure S3K–L), an FG-repeat nucleoporin that contributes to Mex67 docking at the nuclear basket ^84,85^, suggesting age-dependent NPC/basket remodeling that compromised Mex67 docking and mRNA export. In line with an export bottleneck, we observed nuclear accumulation of the mRNA-binding protein Pbp1 and components of the cleavage and polyadenylation machinery (Rna14, Hrp1) in some aged cells (Figure S3H–J), consistent with impaired nuclear RNA export. Faced with this backlog, some aged cells also showed increased expression of factors linked to transcriptional elongation and chromatin transit, including Spt4/5 (DSIF)^86^, Pob3–Spt16 (FACT)^87,88^, and Top2^89^ (Figure S3I, S3M), likely a compensatory—yet potentially maladaptive—response to the unmet cytosolic and organellar metabolic demands stemming from the impaired export of needed transcripts. Beyond these mRNA metabolic changes, age-associated protein aggregation and expression changes of enzymes required for tRNA biogenesis, modification, and aminoacylation likely further altered translation landscape in aged cells (Figure S3N-O). Together, these changes point to a shift from productive translation toward mRNA sequestration and decay, reminiscent of stress-induced translational repression^90^, and may contribute to the age-associated decoupling of mRNA and protein expression observed across species^58,91,92^.

Finally, the ubiquitin–proteasome system (UPS) underwent marked remodeling. While the 20S core appears largely preserved, we observed mislocalization of 19S subunits (Rpn5, Rpn9) from the nucleus to the cytosol in a subset of aged cells, accompanied by reductions in several subunits including Rpn8 and Rpn12 (Figure 2G-H, S3P). Ubiquitin-processing and -conjugating enzymes were similarly perturbed: Ubc4, Ubp10, Ubp14, Ubp16, and Ufd2 were reduced or mislocalized, and the proteasome regulator Ubx4 declined (Figure S3P-Q). Notably, reduction and aggregation of Ubp10 in the nucleolus directly links proteostasis decline to chromatin silencing and rRNA biogenesis: Ubp10 deubiquitinates histone H2B to support Sir2-dependent telomeric silencing and also deubiquitinates Rpa190, stabilizing RNA polymerase I and promoting rRNA synthesis (Figure 2I, S3P) ^93,94^. Functionally akin to the proteasome’s ATP-dependent substrate-handling machinery, Cdc48 (p97/VCP) is a separate ATPase that extracts ubiquitinated substrates and can deliver them for proteasomal processing^95^. Interestingly, we observed a significant increase in the chaperone-ATPase Cdc48 and its spreading from the nucleus into the cytosol (Figure S3P, S3R), likely reflecting an increased cytosolic burden of ubiquitinated or misfolded substrates and compensatory engagement of ATP-driven segregase/extractase activity when 19S-dependent proteasome function is perturbed. However, this compensation was likely not successful, as Otu1, a deubiquitylation enzyme that binds to Cdc48p, relocated from the nucleus to the cytosol and was significantly reduced in many aged cells (Figure S3P, S3S-T). These combined defects in substrate ubiquitylation, chain processing, and proteasome localization suggest that the UPS loses both capacity and spatial organization during aging.

### Age-associated organelle proteostasis defects and organelle-state heterogeneity

Organelle-specific proteostasis systems exhibited widespread decline. Within mitochondria, we observed frequent reduction or aggregation of factors required for respiratory-chain assembly (e.g., cytochrome c oxidase chaperones Sco1/2 and Afg1; bc1 assembly factor Mzm1), mitochondrial proteases (Oma1, Cym1, Mgr1), and components linked to mitochondrial translation and tRNA charging (Mba1, Rsm22, Mrm1, Msw1) (Figure 3A-B, S4A-B). In parallel, aggregation of Phb2 and loss of the import-associated chaperone Zim17 point to reduced folding capacity for newly imported proteins (Figure 3A-B, S4A-B). Notably, the Fe–S biogenesis co-chaperone Jac1 declined early, which may contribute to defects in other mitochondrial proteins dependent on Fe–S as a cofactor (Figure 3A-B, S4A-B, see “mitochondrial dysfunction” below). These proteostasis defects were reinforced by impaired key mitochondrial import machinery: Tim21 was lost early, while many TOM/TIM components (Tom7, Tim44, Tim50) showed increasing aggregation with age (Figure 3B, S4A). Together, these changes likely contribute to mitochondrial biogenesis defects and dysfunction.

**Figure 3.**
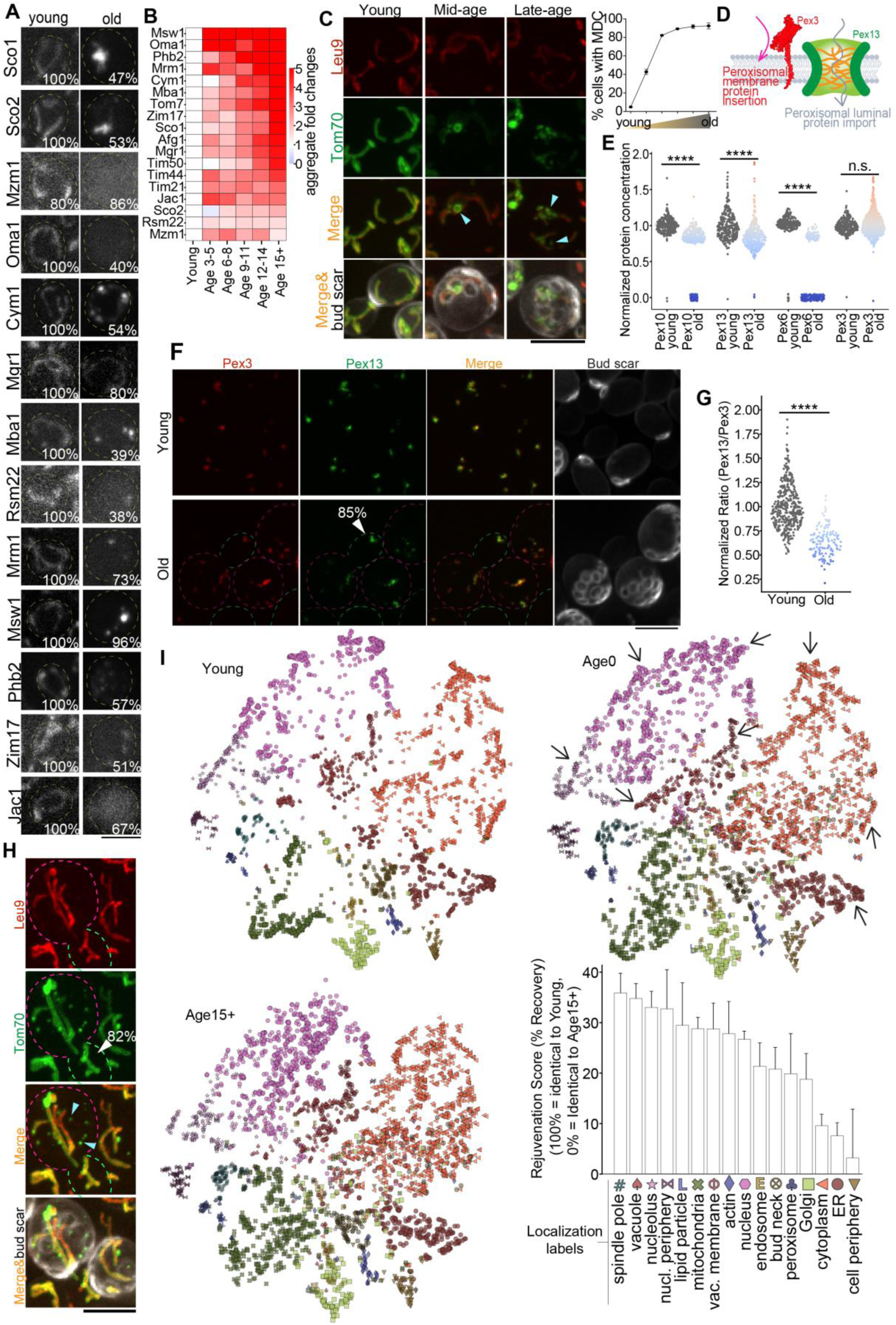
Loss of proteostasis within organelles generate heterogeneous population of organelles. **(A**, **B)** Representative images of different mitochondrial proteostasis machineries (A) and quantification of their age-associated increase of protein aggregation (B). The inset percentage indicates, for each age group, the fraction of cells exhibiting the phenotype shown in the representative images (same definition used in other figures). **(C)** Representative images and quantification of age-associated formation of mitochondria-derived compartments. Cyan arrowheads indicate these mitochondria-derived compartments. Shown are mean and standard error. **(D**, **E)** Schematic illustrating machineries involved in the two pathways of peroxisome protein import (D) and quantification of their age-associated change in protein concentration (E). Each dot represents an individual old cell, with values normalized to the mean protein level in the corresponding young cells. Dot color indicates fold change relative to the young-cell mean (red: increase; blue: decrease; same in other figures). **(F**, **G)** Representative images and quantification showing an age-associated decrease in the Pex13:Pex3 ratio in aging mother cells, followed by restoration of this stoichiometric ratio in rejuvenated daughter cells. Magenta outlines indicate old mother cells, and green outlines indicate young cells. The white arrowhead marks an example of a daughter cell generated by an old mother cell. Among old cells that undergo budding, 85% of daughter cells exhibit this rejuvenation (N=105 old cells). **(H)** Representative images and quantification showing the asymmetric segregation of mitochondria and mitochondria-derived compartments between aged mother cell and the rejuvenating daughter cell. The white arrowhead marks an example of a daughter cell generated by an old mother cell. Among old cells that undergo budding, 82% of daughter cells exhibit this rejuvenation. N= 356 old cells. **(I)** Comparison of proteome-wide localization maps in young cells, aged mother cells (15 plus generations), and newborn daughter cells (age 0). Because we imaged live cells after old-cell purification, purified aged mothers continued dividing and produced newborn age-0 daughter cells. Each dot represents a protein, with dot colors and shapes indicating compartment assignment as shown in the legend below the bar graph. Ensemble-model–quantified localization scores for each protein were plotted in tSNE space and normalized to enable comparisons across ages. Arrows indicate regions showing visible reversal of age-associated dispersion in the localization map in newborn daughters. The inset bar graph shows the calculated rejuvenation score based on differences in localization scores of the same proteins among age-0 daughters, age-15+ mothers, and young cells. A rejuvenation score of 100% indicates a complete return to the young state, whereas 0% indicates a state indistinguishable from age-15+ mother cells. Shown are mean and standard error. Scale bar: 5μm. See table S5 for number of cells quantified.

Importantly, these changes did not reflect a uniform loss of mitochondrial identity; instead, mitochondrial proteostasis failure was modular and heterogeneous. Many mitochondrial proteins retained normal expression in old cells (Figure S4B), whereas specific modules lost solubility, mislocalized, or aggregated (see “mitochondrial dysfunction” below). For example, the outer-membrane import receptors Tom20 and Tom70 progressively aggregated, with a subset incorporated into mitochondria-derived compartments^96^, further weakening import capacity and generating a heterogenous mitochondrial population (Figure 3C, S4C). Consistent with within-cell heterogeneity, mitochondrial matrix proteins such as Ilv1 and Rsm7 displayed partial colocalization and variable enrichment across individual mitochondria in aged cells (Figure S4D).

Similar to mitochondria, peroxisomal import machinery was also destabilized. Most luminal proteins of the peroxisome are imported from the cytosol by the soluble receptor Pex5 via the translocation pore Pex13 into the lumen of the peroxisome, whereas peroxisomal membrane proteins are delivered by Pex19 and inserted via Pex3-dependent pathways^97–99^ (Figure 3D). Pex5 is then recycled back to the cytosol by the ATPase Pex1/6 via a retrotranslocation channel formed by Pex10^100^. During aging, Pex3 remained comparatively stable, whereas Pex10, Pex13, and Pex6 declined markedly (Figure 3E, S4E), a pattern predicted to compromise peroxisomal biogenesis and function. These shifts altered the relative abundance of import machinery components compared with Pex3 (Figure 3F–G, S4F–G). Interestingly, aged cells displayed a mosaic peroxisomal population in which individual peroxisomes varied substantially in their Pex10:Pex3 ratios (Figure S4F, S4H), supporting the idea that organelle proteostasis failure is accompanied by within-cell organelle-state diversity.

Finally, these age-associated organelle proteostasis defects were partially reset during asymmetric cell division. Daughters of aged mothers inherited mitochondria largely free of Tom20/Tom70 aggregates and mitochondria-derived compartments, which were preferentially retained in the mother (Figure 3H, S4C). Likewise, the age-dependent reduction in Pex13 was restored in daughters produced by aged mother cells (Figure 3F). Extending beyond these examples, analysis of newly born age-0 cells within our purified old-cells revealed a broader, proteome-wide trend toward restored localization and morphology across many compartments (Figure 3I, Video S1, Table S2). Together, these observations support a model in which replicative aging drives a breakdown of organellar proteostasis that fragments organelle states within a cell, and that asymmetric division counteracts this erosion by re-establishing a more youthful organelle proteome in the newborn daughter.

### Dysregulated nutrient signaling and remodeling of nutrient transporters during aging

Aging is accompanied by rewiring of conserved nutrient-sensing pathways that normally balance anabolic growth with stress defenses (e.g., TORC1, sirtuins; IIS in animals)^2^. Despite constant nutrient-replete conditions during old cell enrichment, single-cell analysis reveals heterogenous, compartment-specific changes that point to dysregulated nutrient sensing and response programs that are decoupled from extracellular nutrient availability.

#### Amino acid signaling

In line with tissue-specific reports that mTOR activity can increase with age^101^, we observed increased vacuolar enrichment of the Rag GTPase module Gtr1–Gtr2, a key upstream activator of TORC1(Figure 4A)^102^. Because TORC1 signaling integrates amino acid and nitrogen availability^102–105^, we next examined nutrient transporter remodeling. Multiple amino-acid permeases (Dip5, Bap2, Agp1, Lyp1, Tat1) were depleted from the plasma membrane and accumulated in the vacuole, consistent with endocytic downregulation (Figure 4B, S5A). This pattern suggests reduced capacity for amino-acid import, alongside a relative shift toward ammonium uptake, as Mep1/2 increased (Figure S5B). One possible driver is an age-associated rise in intracellular amino acids (e.g., cysteine), as recently reported^106^. Notably, Met4, a central regulator of sulfur amino-acid biosynthesis, remained unchanged rather than showing the expected sulfur-replete inactivation pattern^107^, raising the possibility of defective sulfur-metabolite sensing in which elevated intracellular sulfur amino acids fail to engage Met4 feedback control in aged cells (Figure S5C).

**Figure 4.**
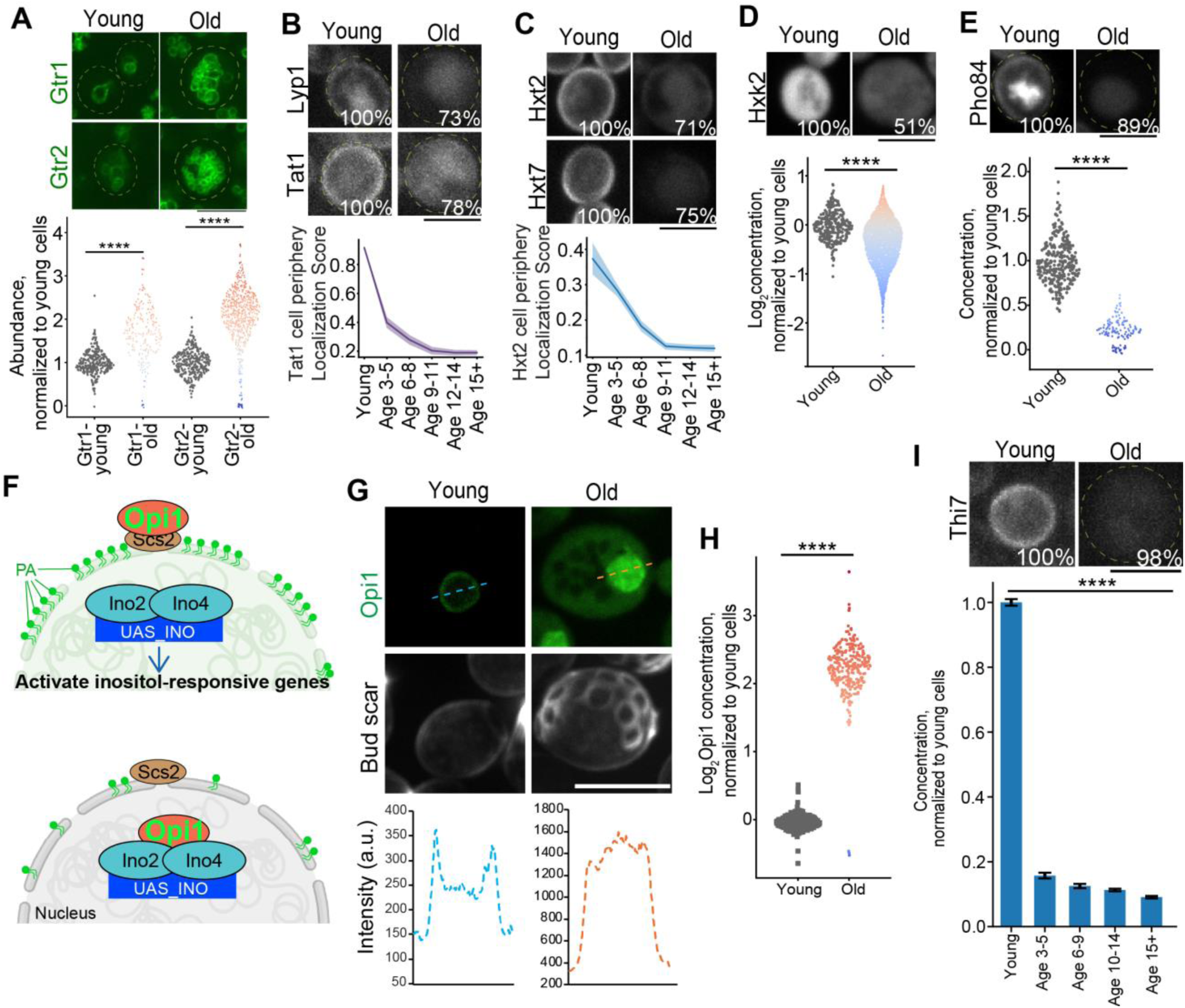
Age-associated molecular changes underlying deregulated nutrient sensing. **(A)** Example and quantification of protein abundance for Gtr1/2 in young and old cells. Each dot represents an individual old cell, with values normalized to the mean value of the corresponding protein in young cells (same for other figures). Dot color indicates fold change relative to the young-cell mean (red: increase; blue: decrease; same in other figures) **(B**, **C)** Representative images and quantification of cell periphery localization and expression changes of amino acid transporters (B) and glucose transporters (C) in young and old cells. Shown are mean and standard error (shade). The inset percentage indicates, for each age group, the fraction of cells exhibiting the phenotype shown in the representative images (same definition used in other figures). **(D**, **E)** Representative images and quantification of Hxk2 (D) and Pho84 (E) in young and old cells. **(F**-**H)** Schematic illustrating of UAS_INO gene regulatory mechanism (F) and Representative images and quantification of Opi1 localization (G) and expression (H) in young and old cells. **(I)** Representative images and quantification of Thi7 in young and old cells. Scale bar: 5μm. See table S5 for number of cells quantified.

#### Glucose sensing and glycolytic input

Beyond amino acids, we observed reduced expression and frequent intracellular retention of glucose transporters Hxt2 and Hxt7 away from the plasma membrane (Figure 4C). Hexokinase Hxk2, which was uniformly high in young cells, declined and showed increased cell-to-cell variability with age (Figure 4D), suggesting reduced glucose phosphorylation capacity and more heterogenous glycolytic activity in aged cells. In addition to glucose phosphorylation, hexokinases can participate in glucose sensing; for example, Hxk2’s mammalian homolog GCK/HK4 serves as a glucose sensor in pancreatic cells and contributes to insulin secretion^108,109^. Together, reduced surface availability of glucose transporters and diminished Hxk2 suggest impaired glucose sensing and attenuated glucose-responsive signaling in aged cells, despite periodic media refresh (every 2 h) to maintain nutrient-replete conditions. In line with weakened glucose-responsive regulation, many aged cells showed increased expression and aggregation of Adh3 (see “mitochondrial dysfunction”, Figure 5E, S6H), which is normally glucose-repressed^110^.

**Figure 5.**
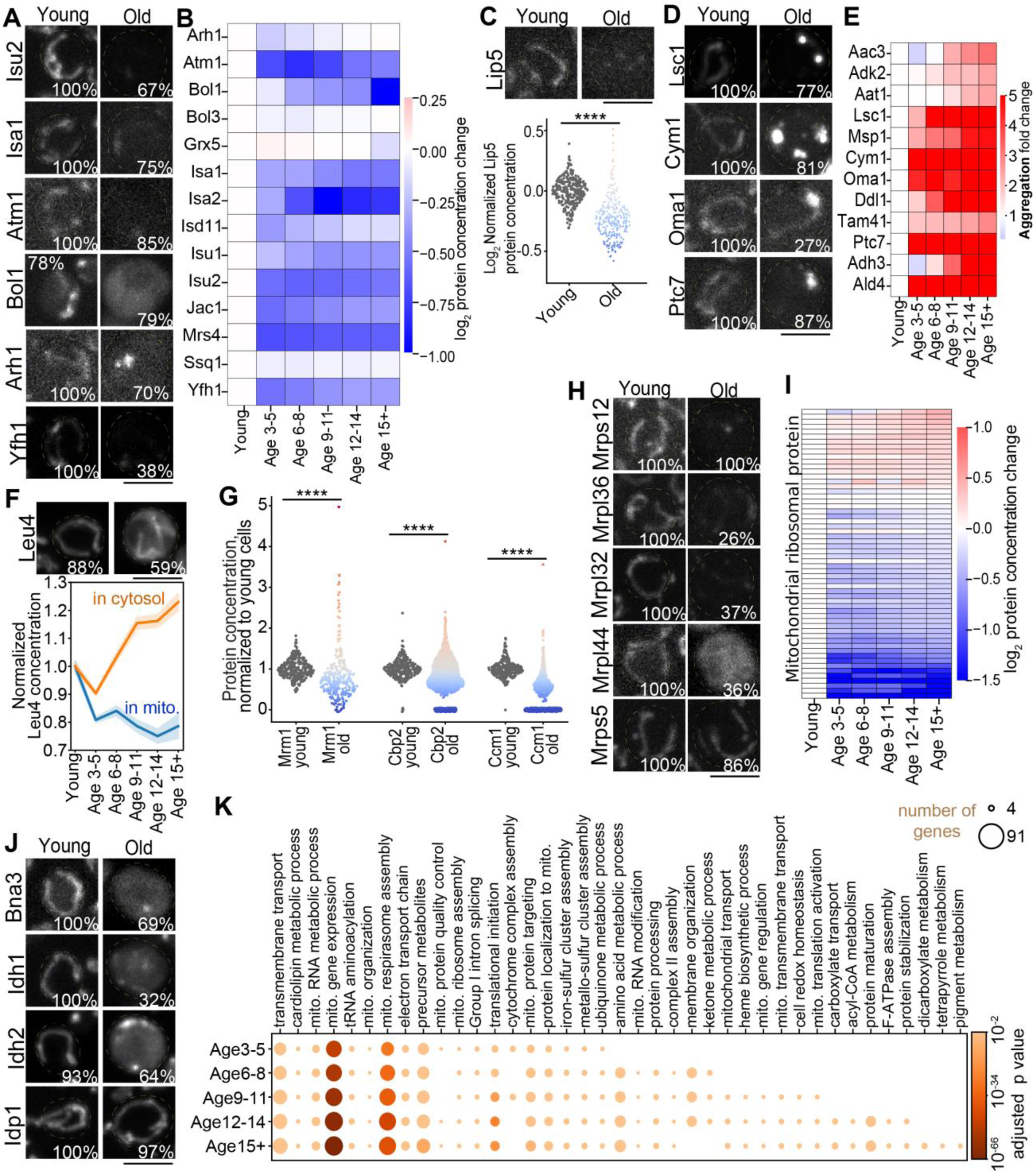
Age-associated molecular changes underlying dysfunctional mitochondria. **(A)** Representative images and quantification of protein concentrations for different ISC enzymes in young and old cells. The inset percentage indicates, for each age group, the fraction of cells exhibiting the phenotype shown in the representative images (same definition used in other figures). **(B)** Quantification of protein concentrations for components of ISC pathway during aging. **(C)** Representative images and quantification of protein concentrations for Lip5 in young and old cells. Each dot represents an individual old cell, with values normalized to the mean protein level in the corresponding young cells (same for other panels). Dot color indicates fold change relative to the young-cell mean (red: increase; blue: decrease; same in other figures). **(D-E)** Representative images and quantification of protein aggregation for different mitochondrial metabolic enzymes and structural proteins in young and old cells. **(F)** Representative images and quantification of Leu4 protein concentration in the cytosol and mitochondria of young and old cells. Shown are mean and standard error (shade). **(G)** Quantification of protein concentrations for Mrm1, Cbp2, and Ccm1 in young and old cells. **(H-I)** Representative images and quantification of mitochondrial ribosomal protein concentrations in young and old cells. Not all mitoribosomal proteins changed with age; for example, Mrps5 remained relatively stable during aging. **(H)** Representative images and quantification of subcellular localization changes for NAD^+^-metabolism related proteins in young and old cells. **(I)** GO term biological process enrichment for the mitochondrial proteins showing concentration changes or protein aggregation during aging. Scale bar: 5μm. See table S5 for number of cells quantified.

#### Collapse of phosphate sensing and transporter

A less-appreciated axis of dysregulation is phosphate (Pi) homeostasis. The high-affinity H⁺/Pi symporter Pho84 is the principal phosphate “transceptor” in yeast that senses ambient Pi and transports it to the cytosol, coupling phosphate import to signaling outputs through PKA and TORC1^111–113^. We found that Pho84 was lost from the plasma membrane during aging, coincident with a marked decline of Pho86, the ER chaperone required for Pho84 trafficking to the surface (Figure 4E, S5D)^111,114,115^. In parallel, the auxiliary Na⁺/Pi transporter Pho89 was reduced (Figure S5E), and Spl2, a regulator that helps enforce appropriate phosphate-transporter usage, was strongly diminished (Figure S5F). The coordinated loss of Pho84, Pho86, Pho89, and Spl2 points to broad erosion of the PHO network, with predicted consequences for Pi uptake and Pi-dependent signaling during aging.

#### Phospholipid/inositol transcriptional control

A second, previously underappreciated axis of aging involves defects in phospholipid and inositol transcriptional control. In young cells, Ino2–Ino4 activates the transcription of the UAS_INO gene network and inositol-responsive genes, while Opi1 is sequestered at the perinuclear ER by phosphatidic acid (PA) and Scs2; when inositol is abundant, PA levels drop as PA is converted to phosphatidylinositol (PI), allowing Opi1 to enter the nucleus and repress UAS_INO targets (Figure 4F)^116,117^. During aging, we observed selective loss of Ino2 together with increased Opi1 abundance and a shift from the ER to the nucleus (Figure 4G–H, S5G)—a coherent signature of sustained UAS_INO repression. In line with this, Scs2, an ER anchor that restrains Opi1^117^, exhibited pronounced age-dependent localization changes (Figure S5H), potentially facilitating Opi1 release and nuclear accumulation. In parallel, Faa3, which enables utilization of exogenous fatty acids for phospholipid synthesis, was progressively lost in aged cells (Figure S5I). Such Faa3 loss may contribute to altered phospholipid metabolism and PA-dependent Opi1 regulation. Together, these changes support dysregulation of membrane-lipid biosynthesis and ER homeostasis.

#### Micronutrient logistics

In addition to macronutrient dysregulation, aging cells also exhibited widespread alterations in micronutrient homeostasis (e.g., metal ions and vitamin-derived cofactors). Because these cofactors are integral to mitochondrial respiration, redox buffering, and cofactor-dependent catalysis, even modest imbalance can propagate into broader metabolic dysfunction^118–120^. We observed that the copper importer Ctr1 was markedly upregulated, while Fre1, a cell-surface ferric/cupric reductase that supplies Fe³⁺/Cu²⁺ to their respective transporters, was also elevated (Figure S5J-K). At the vacuole, the Zn²⁺ importer Cot1 increased in ∼half of aged cells, while the TRP-like vacuolar Ca²⁺ channel Yvc1 (TRPY1) frequently aggregated (Figure S5J), consistent with disrupted ion storage and vacuolar stress. Beyond metals, the thiamine (vitamin B₁) transporter Thi7 declined markedly with age (Figure 4I). Because thiamine is converted to thiamine pyrophosphate, an essential cofactor for central metabolic enzymes—including mitochondrial pyruvate dehydrogenase and α-ketoglutarate dehydrogenase, as well as cytosolic transketolase^121^—reduced Thi7 could lead to an emerging cofactor limitation that would exacerbate mitochondrial and metabolic dysfunction.

Together, in the context of nutrient-replete culture conditions during our old-cell enrichment, these single-cell signatures support a model in which aging decouples extracellular nutrient availability from intracellular nutrient-sensing states, producing heterogeneous, compartment-specific remodeling of transport, signaling, and transcriptional control programs across multiple nutrient homeostasis pathways.

### Mitochondrial dysfunction in metabolism and organelle homeostasis

Our proteome-wide microscopy revealed a broad spectrum of mitochondrial dysfunction during replicative aging. The dominant signature is not a uniform loss of mitochondrial proteins, but modular failure across cofactor biogenesis, mitochondrial gene expression/translation, membrane/lipid homeostasis, and protein quality control, accompanied by widespread protein aggregation and selective mis-localization. Together, these changes indicate a multifaceted mitochondrial decline that is predicted to compromise respiratory competence, metabolic flexibility, and organelle quality control.

#### Iron–sulfur cluster (ISC) biogenesis and cofactor synthesis

Fe–S cluster assembly emerged as one of the earliest and most prominent pathways disrupted during aging (Figure 5A-B). The early scaffold protein Isu2^122^, required for 2Fe–2S cluster formation, was lost in the majority of cells at early stage of aging, whereas the late-acting Isa1, which mediates 4Fe–4S cluster maturation^122^, declined more gradually (Figure 5A-B). Downstream components of Fe–S synthesis—including the Fe-S-insertion chaperone Jac1, the mitochondrial Fe-S exporter Atm1, and Bol1, which mediates 4Fe–4S cluster transfer to specific client proteins^122^—either decreased in abundance or mislocalized to the cytosol in aged cells (Figure 5A-B, S6A). In contrast, the related mitochondrial BolA-like factor Bol3 remained relatively stable (Figure 5B, S6B). Additional ISC-linked assembly factors (e.g., Arh1, Yfh1) showed aggregation or reduction (Figure 5A-B, S6C). Collectively, these changes point to a systemic collapse of Fe–S biogenesis, likely destabilizing Fe–S–dependent proteins/pathways, many of which declined in abundance and/or formed aggregates (Figure S6D-E).

One such example is the lipoic acid synthesis pathway. Bol1 facilitates the incorporation of Fe–S clusters into some mitochondrial client proteins, including the lipoate synthase Lip5^123^. In aged cells, Lip5 was depleted in more than half of the population (Figure 5C), coincident with reduction/aggregation of lipoate-dependent subunits (Lat1, Kgd2, Gcv3) of pyruvate dehydrogenase, α-ketoglutarate dehydrogenase, and glycine cleavage system, along with their associated proteins (Figure S6F–G)^124^. Interestingly, a similar decline in lipoic acid biogenesis has been reported during animal and human aging due to the reduction of Lip5 ortholog LIAS, and lipoic acid supplementation has been shown to preserve stem-cell function by sustaining mitochondrial TCA-cycle enzymes and PGC-1α activity and supporting telomere integrity^125–128^.

#### Mitochondrial protein aggregation

Many mitochondrial enzymes and structural proteins progressively formed puncta or insoluble assemblies, spanning metabolite transport, TCA-cycle metabolism, and protease/quality-control systems (e.g., Aac3, Adk2, Aat1, Lsc1, Msp1, Cym1, and Oma1) (Figure 5D-E, S6H). Aggregation was especially pronounced among proteins critical for ubiquinone and cardiolipin/phospholipid metabolism—Ddl1, Tam41, and Ptc7—consistent with impaired lipid remodeling and cristae maintenance (Figure 5D-E, S6H). Several enzymes showed conspicuous age-dependent remodeling: Adh3, normally repressed by glucose^110^, was strongly upregulated and formed bright aggregates, while Ald4 formed fibers in ∼20% of aged cells—resembling its filamentation under sporulation and nutrient-limiting conditions^129^ (Figure 5E, S6H). Because these aging cells were maintained in nutrient-replete YPD, these starvation-like phenotypes point to deregulated nutrient-sensing, another hallmark of aging, with a subset of cells engaging a partial, heterogeneous starvation-response program despite abundant extracellular nutrients.

#### Mis-localization of mitochondrial enzymes to the cytosol

Cytosolic relocalization of mitochondrial proteins was another recurring lesion. Leu4, the main isozyme for leucine biosynthesis, shifted to the cytosol in aged cells (Figure 5F). Because translation from a second in-frame AUG can generate a cytosolic Leu4 isoform^130,131^, this pattern may reflect age-dependent changes in translation initiation or isoform usage. Several additional mitochondrial enzymes showed similar mislocalization, including Fum1 (fumarase), Aim45, Cir1 (a Yfh1-interacting protein; ortholog of mammalian electron transfer flavoprotein subunit ETF-beta), and the glycine cleavage component Gcv1 (Figure S6I). Together, these patterns indicate that mitochondrial dysfunction during aging arises not only from protein depletion and aggregation, but also from failure to maintain proper mitochondrial targeting and retention of key enzymes.

#### Collapse of mitochondrial translation and ribosome integrity

Impaired mitochondrial translation is a conserved feature of mitochondrial aging^132,133^. Splicing and RNA-processing factors—including Cbp2 (COB intron splicing), Ccm1 (15S rRNA stabilization), and Mrm1 (21S rRNA methyltransferase)—were depleted with Mrm1 was absent in ∼70% of cells by mid-age (Figure 5G, S6J). Numerous mitoribosomal proteins (e.g., Mrpl8, Mrpl32, Mrpl36, Mrpl40, Mrps12, Mrps35, Rml2/uL2m, Rsm28/mS46, Img1/bL19) declined or mislocalized during aging (Figure 5H-I). Notably, Mrpl44, a mitoribosomal large subunit protein, mislocalized from mitochondria to the cytosol, highlighting a breakdown in mitoribosome integrity (Figure 5H). Translation fidelity and mt-tRNA charging capacity also eroded: Her2, a component of the GatFAB amidotransferase complex that prevents mistranslation of Gln codons^134^, and multiple aminoacyl-tRNA synthetases (Mst1, Msw1) aggregated in aged cells (Figure S6K, S6L). In parallel, several factors (e.g., Atp25, Pet54, Pet100; Mdm38, Mzm1, Cbp4, Afg1) that coordinate mitoribosome-translated nascent subunit production and their incorporation into OXPHOS complexes were lost or aggregated (Figure S4A, S6K-M), consistent with a profound failure of OXPHOS biogenesis.

#### Metabolic rewiring and cofactor depletion

Beyond structural breakdown, aged mitochondria showed selective metabolic rewiring through redistribution of key enzymes between mitochondria and the cytosol. Bna3 (kynurenine aminotransferase) relocalized from mitochondria to the cytosol in aged cells (Figure 5J), which may siphon kynurenine into the kynurenic-acid branch in the cytosol at the expense of kynurenine flux into the mitochondria for *de novo* NAD⁺ synthesis, potentially contributing to age-associated NAD⁺ decline^135,136^. In parallel, the NAD⁺-dependent isocitrate dehydrogenases Idh1/2 mislocalized from mitochondria to the cytosol in many aged cells, while the NADP⁺-dependent isoform Idp1 remained mitochondria-restricted, suggesting a selective breakdown of mitochondrial NAD⁺-coupled TCA metabolism (Figure 5J)^137^. Fum1’s mislocalization to the cytosol and the gradual reduction of Cit1 (citrate synthase) further compromised TCA cycle integrity (Figure S6I, S6N-O).

Cofactor biosynthesis was also affected. Cat2, a carnitine acetyl-CoA transferase that converts acetyl-CoA to acetylcarnitine for membrane transport, relocalized from cytosol to mitochondria in some aged cells (Figure S6P). Ach1, the mitochondrial acetyl-CoA hydrolase, accumulated as puncta after mid-age (Figure S6P). Hem15, the terminal heme biosynthetic enzyme, declined in over half of aged cells while early heme steps (e.g., Hem1) remained intact (Figure S6Q-R). Consistent with broader cofactor/precursor imbalance, multiple cofactor-dependent metabolic enzymes, including those involved in amino-acid metabolism and transport (Ilv1/2, Gcv1/3, Arg8, Odc2), were reduced or aggregated (Figure S6Q-R). Together, these alterations highlight disrupted cofactor homeostasis and metabolic wiring in aging mitochondria.

#### A stepwise trajectory of mitochondrial failure

Integrating these phenotypes revealed a staged progression of mitochondrial decline rather than a stochastic collapse (Figure 5K). Early changes were enriched for mitochondrial RNA metabolism, mitoribosome biogenesis, and cofactor/precursor pathways (notably Fe–S cluster biogenesis). These were followed by defects in membrane organization and protein import, alongside disruptions in amino-acid metabolism. Later stages showed impaired heme biosynthesis, redox homeostasis, and acyl-CoA metabolism, culminating in defective F1F0-ATP synthase assembly and broad metabolic remodeling. Consistent with this staged model, many mitochondrial proteins remained relatively stable until late life, including Idp1, Hem1, and Coa1/Cor1 (cytochrome c oxidase and bc1 assembly factors) (Figure S6S).

### Nucleolar Dysfunction and Ribosome Biogenesis Defects

Nucleolar dysfunction has emerged as a conserved feature of aging, reflected by nucleolar enlargement across species and by the strong impact of translational fidelity on lifespan^12,79–81^. Mutations that increase fidelity extend lifespan in yeast, worms, and flies, whereas disruption of ribosome biogenesis factors—including SBDS (yeast Sdo1) and Tif6—causes ribosomopathies, impaired translation, and reduced organismal fitness^138,139^. Our single-cell atlas extended this framework by revealing a spatial failure mode of ribosome biogenesis during aging: the nucleolus progressively lost its identity as a compartmentalized assembly hub. Ribosome biogenesis is a multistep pathway centered in the nucleolus, encompassing rRNA transcription, endonucleolytic cleavage, chemical modification, and hierarchical assembly of preribosomal particles (Figure 6A). Rather than a uniform decline in ribosome-biogenesis protein abundance, we observed a widespread loss of nucleolar confinement during aging, with many processing, remodeling, and quality-control factors escaping into the nucleoplasm and frequently forming aggregates. This spatial breakdown is predicted to uncouple sequential maturation steps and erode both efficiency and fidelity of ribosome production.

**Figure 6.**
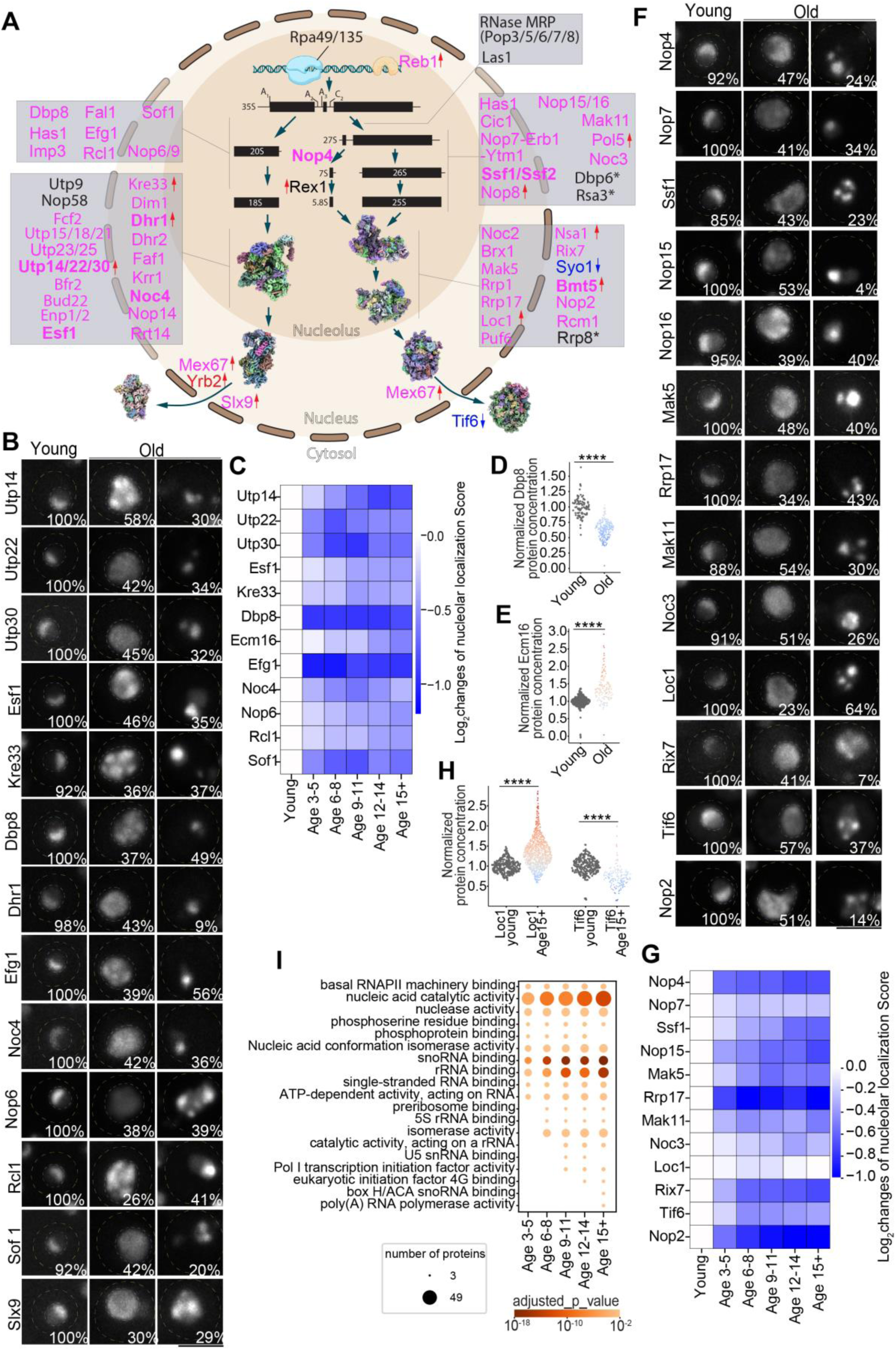
Age-associated changes in nucleolus and ribosome biogenesis. **(A)** Schematic of major biogenesis steps for the small (SSU) and large (LSU) ribosomal subunits. Key assembly factors and rRNA processing/modification enzymes are annotated at each step. Proteins that relocalized from the nucleolus to the nucleoplasm during aging are shown in pink, whereas proteins whose expression decreased with age are shown in blue; proteins without detectable localization or expression changes are shown in black. Blue downward and red upward arrows denote age-associated down- and up-regulation, respectively. * indicates proteins that aggregate with age. Ribosomal subunit models were generated from PDB entries 6G4W, 6G5H, and 6ZQF for SSU and 3J7P, 3JCT, 6C0F, and 6LSR for LSU. **(B-C)** Representative images (B) and quantification of nucleolar localization (C) for factors involved in SSU maturation in young and old cells. The inset percentage in (B) indicates, for each age group, the fraction of cells exhibiting the phenotype shown in the representative images (same definition used in other figures). For old cells, examples of both nucleolar escape and aggregation are shown in (B). See Figure S7F-G for additional proteins. **(D-E)** Quantification of protein concentrations for Dbp8 (D) and Ecm16 (E) in young and old cells. Each dot represents an individual old cell, with values normalized to the mean protein level in the corresponding young cells (same for other panels). Dot color indicates fold change relative to the young-cell mean (red: increase; blue: decrease; same in other figures). **(F-G)** Representative images (F) and quantification of nucleolar localization (G) for factors involved in LSU maturation in young and old cells. For old cells, examples of both nucleolar escape and aggregation are shown in (F). See Figure S8A-B for additional proteins. **(H)** Quantification of protein concentrations for Loc1 and Tif6 in young and old cells. **(I)** GO term molecular function enrichment for the nucleolar proteins showing concentration changes or protein aggregation during aging. Scale bar: 5μm. See table S5 for number of cells quantified.

#### Transcription persists, but coordination begins to drift

Core Pol I transcription components—including Rpa49 and Rpa135 (Pol I subunits) and the scaffold Rrp5—maintained nucleolar localization until late in lifespan (Figure 6A, S7A–C), suggesting that rRNA transcription remained broadly intact through much of aging. Several early cleavage machineries (e.g., Nop58 and RNase MRP subunits) were also comparatively stable (Figure 6A, S7A–D). In contrast, transcriptional regulation and its coupling to early pre-rRNA processing diverged earlier: Reb1 increased in a subset of aged cells (Figure 6A, S7E), and the helicase Dbp3, required for A3 cleavage of nascent 35S pre-rRNA^140^, redistributed from the nucleolus into the nucleoplasm (Figure S7E). This spatial separation is expected to impair A3-site processing despite sustained Pol I transcription.

#### Small subunit (SSU) processome disorganization

A dominant early lesion was the breakdown of the SSU processome, which normally organizes pre-18S folding, cleavage, and modification in a tightly choreographed nucleolar environment^141^. In aged cells, we observed coordinated nucleolar escape across multiple SSU modules: UTP/U3-linked constituents and associated SSU factors redistributed into the nucleoplasm and frequently formed aggregates (Figure 6A–C, S7F–I). The broad loss of nucleolar confinement across SSU components is consistent with a shared underlying defect in nucleolar retention that would be expected to compromise ordered pre-18S maturation.

Within this broader failure, several node proteins provide mechanistic anchors. The 18S rRNA acetyltransferase Kre33 became abnormally elevated and aggregated in a subset of aged cells (Figure 6B, S7J) ^142,143^. Notably, Kre33’s human homolog NAT10 is implicated in progeroid and physiological aging phenotypes, and NAT10 inhibition rescues cellular defects in progeroid patient cells and aged human donor cells, and improves healthspan in progeroid mouse models ^144–146^. This cross-species parallel highlights that remodeling enzymes acting on early SSU intermediates can be particularly consequential nodes.

A second signature within this SSU assembly defect is the failure of ATP-dependent rRNA remodeling. The DEAD-box helicases Dbp8 and Has1 (early rRNA folding/cleavage) became diffuse throughout the nucleus or reduced (Figure 6A-D, S7F-I) ^147^, while Ecm16/Dhr1, which releases U3 snoRNA to allow central pseudoknot formation^148^, increased in expression level yet lost nucleolar localization (Figure 6A-C, 6E). Additional SSU processome factors that support particle integrity (e.g., Efg1, Fal1, Faf1) showed similar nucleolar escape and aggregation (Figure 6A-C, S7F-I) ^141,147^. Conceptually, these changes convert an ordered remodeling pipeline into a mislocalized, aggregation-prone state—precisely the kind of failure expected to generate aberrant intermediates and delay or derail 40S maturation.

Finally, architectural scaffolds that coordinate domain folding and cleavage also lost nucleolar confinement. Early 40S organizers—including Imp3, Krr1, and the Nop14–Noc4–Enp1 scaffold that engages the pre-18S 3′ domain—redistributed into the nucleoplasm and frequently aggregated (Figure 6A-C, S7F-I) ^141^. Additional nucleolar RNA-binding factors (Nop6, Nop9) followed the same pattern, and the A2 cleavage factor Rcl1 was likewise displaced (Figure 6A-C, S7F-I) ^141^. Mislocalization of U3-associated factors (e.g., Rrt14, Sof1) further compounded the spatial uncoupling of cleavage and folding steps that normally occur in the nucleolus (Figure 6A-C, S7F-I)^141^.

Along with widespread upstream defects, we observed an increase in Yrb2 and Slx9, factors involved in export of late pre-40S particles, potentially reflecting compensatory upregulation of downstream steps in response to defective 40S assembly (Fig. 6A–B, S7K–L).

#### LSU assembly failure centers on an ITS2 weak point

Large-subunit assembly followed a similar pattern: aging preferentially destabilized nucleolar modules that scaffold and protect early pre-60S architecture^141^. A key signature was the fragility of the ITS2-centered network. Nop4, required for ITS1 processing and early LSU maturation^149^, became mislocalized and aggregated in most aged cells (Figure 6A, 6F-G). Factors that stabilize the ITS2 region—including Brx1, the Nop7–Erb1–Ytm1 subcomplex, Nop15, and Nop16—frequently lost nucleolar confinement and formed aggregates (Figure 6A, 6F-G, S8A-C) ^141^. Cic1/Nsa3, which binds the ITS2 region and supports early nucleolar particle architecture^141^, similarly became mislocalized and aggregated (Figure 6A, S8A-C), reinforcing the idea that ITS2-centered intermediates are particularly vulnerable during aging. These changes occurred alongside mislocalization of Ssf1/Ssf2, which normally mark the nucleolar 27S A2 state and shield ITS2 from premature processing (Figure 6A, 6F-G, S8A-B)^141^. Together, these changes likely compromise the ITS2-centered stabilization network, eroding the scaffold that normally supports ordered remodeling and progression.

Upstream of this ITS2 module, the Nop8–Rsa3–Dbp6 network—critical for early pre-rRNA compaction steps^141^—showed early reduction and aggregation (Figure 6A, S8A-C), consistent with early-onset defects in LSU architecture that likely ripple into ITS2 processing. Additional remodeling enzymes and scaffolds further eroded LSU progression: DEAD-box helicase Mak5 became diffuse throughout the nucleus (Figure 6A, 6F–G)^141^; Rrp17 aggregated (Figure 6A, 6F–G)^141^; and nucleolar scaffolds such as Mak11, Noc3, and Pol5 failed to remain localized (Figure 6F–G, S8A–B), compounding the collapse of early assembly intermediates ^141^.

Late-stage licensing and export checkpoints also weakened in characteristic ways. Loc1 and Puf6, which coordinate 27S processing and chaperone delivery of Rpl43, dispersed into the nucleoplasm and aggregated (Figure 6F–G, S8A–C)^141^. Interestingly, Loc1 expression level increased during aging, likely compensating for defects at this stage of 60S maturation (Figure 6H). Nsa1, which sits on the solvent-exposed side of nucleolar pre-60S particles and helps stabilize the junction of 25S rRNA domains I and II, increased in some aged cells but dispersed, while Rix7, the AAA-ATPase responsible for removing Nsa1 as maturation proceeds, also became mislocalized (Figure 6F-G, S8D)^141^. Downstream, Syo1—the specialized import adaptor for Rpl11—was reduced (Figure S8D), likely contributing to the reduction of Rpl11 in some aged cells (Figure S8E). Finally, Tif6—whose release by the Efl1–Sdo1/SBDS axis licenses productive 60S joining^141^—declined sharply (Figure 6F–H), indicating erosion of a key late quality-control checkpoint.

#### rRNA modification and surveillance enzymes lose nucleolar confinement

Aging disrupted not only the structural assembly factors but also the enzymes that chemically mature and proofread rRNA. These enzymes normally operate in a tightly compartmentalized nucleolar environment in young cells to ensure efficient modification, processing, and surveillance^141^. Dim1 and Bmt5, both rRNA methyltransferases, relocalized from the nucleolus to the nucleoplasm in aged cells and formed aggregates (Figure S7F, S8F). Additional methyltransferases—including Nop2 and Rcm1—became diffuse across the nucleus (Figure 6F-G, S8F-G), and Rrp8 aggregated in ∼30–40% of aged cells (Figure S8F). These changes are poised to reshape rRNA modification patterns and weaken the quality-control checkpoints they support. Moreover, loss of nucleolar confinement of these modification enzymes may increase their access to noncanonical substrates, as shown for the rRNA methyltransferase Spb1, which can also methylate subsets of mRNAs under nutrient limitation^150^.

Surveillance capacity also declined. Rex1/Rnh70, a 3 ′– 5 ′ exonuclease implicated in maturation of 5S/5.8S rRNA and tRNA-Arg3^151^, showed increased expression and greater cell-to-cell variability in some aged cells (Figure S8H). In contrast, the exosome core subunit Rrp43 was reduced in a subset of cells (Figure S8H), likely impairing the clearance of aberrant pre-rRNAs.

Overall, these results identify protein mislocalization and aggregation as primary lesions of nucleolar aging and ribosome biogenesis failure. Notably, these defects unfold in a staged, asynchronous manner across factors and pathways (Figure 6I). Because ribosome biogenesis dominates biosynthetic investment— mRNAs encoding ribosomal proteins comprise ∼50% of Pol II output, and ribosomal proteins account for ∼30% of total proteins^152,153^—early disruption of ribosome assembly is expected to undermine proteostasis. Moreover, any aberrant ribosomes, even, at low abundance, produced under these conditions could further amplify proteotoxic stress, given that reduced translational fidelity is itself a major source of protein damage^154,155^. Thus, by eroding nucleolar organization and spatial coupling of biogenesis steps, aging cells derail ribosome production at multiple levels, providing a direct mechanistic bridge from nucleolar disorganization to proteostasis collapse.

### Overview and cross-species conservation of hallmark-linked proteome remodeling

Our spatial single-cell atlas reveals hundreds of previously unappreciated, yet interconnected age-associated changes in protein concentration, localization, and aggregation that collectively map onto multiple conserved hallmarks of aging as defined in the López-Otín framework (Figure 7A)^1,2^. Beyond cataloging these changes, the atlas provides additional mechanistic anchors for previously reported aging outputs—such as mtDNA instability, increased cryptic transcription, and decoupling between mRNA and protein levels—while uncovering extensive subcellular and spatiotemporal remodeling that is not readily captured by bulk proteomics. Prominent examples include age-dependent loss of compartmental confinement among proteins involved in ribosome biogenesis, mitochondrial biogenesis/metabolism, and nucleocytoplasmic mRNA transport, alongside widespread protein aggregation enriched for mitochondrial and ribosome-biogenesis pathways.

**Figure 7.**
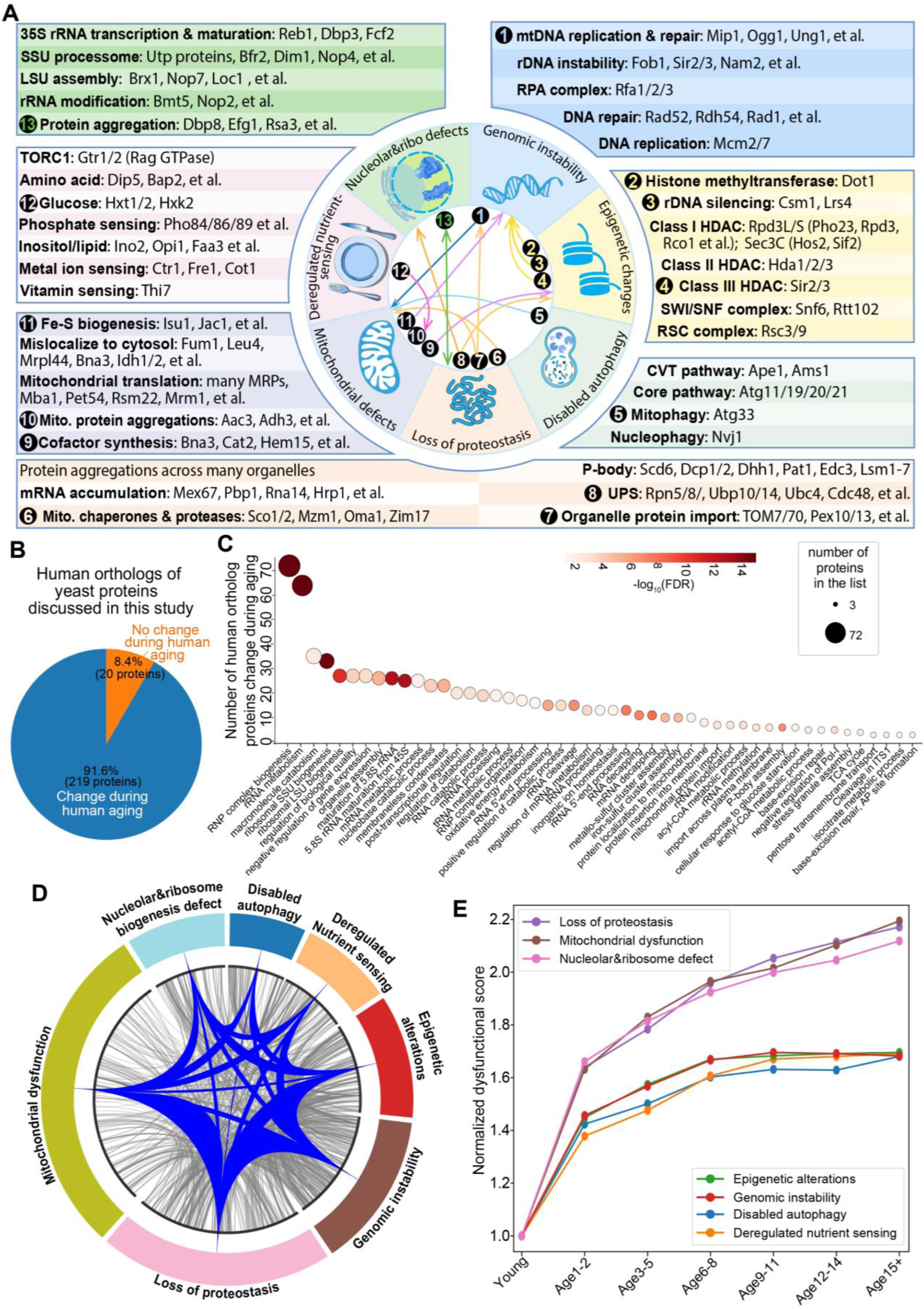
Summary of interconnections and temporal relationship among hallmarks of aging. **(A)** Summary of the molecular changes underlying conserved hallmarks of aging discussed in this study. The central inset depicts functional connections among the hallmark-associated changes highlighted here. For example, organelle import and proteostasis defect (7) may contribute to the selective loss of DNA repair enzymes from mitochondria, which could promote mtDNA instability (1) and, in turn, exacerbate mitochondrial dysfunction. **(B)** Proportion of human orthologs of the hallmark-linked yeast proteins discussed in this study that show age-associated changes in human aging (91.6%, 219 proteins) versus no significant change (8.4%, 20 proteins), based on the human aging proteomics dataset in this recent study^159^. **(C)** Biological process enrichment among the 219 human orthologs that show age-associated changes during human aging. As in yeast, terms related to ribosome biogenesis, mitochondrial function (e.g., Fe–S cluster assembly and the TCA cycle), and P-body assembly are prominently enriched. **(D)** Proteome-scale map of inter-hallmark connectivity. Proteins assigned to each hallmark category are arranged on the outer ring. Chords connect proteins from different hallmark categories when they are functionally linked (see Methods), revealing dense cross-hallmark coupling across the hallmark network. Blue ribbons aggregate the underlying protein–protein links for each hallmark pair (gray chords show individual connections). **(E)** Hallmark-level temporal trajectories during replicative aging. For each age bin, we computed a normalized hallmark dysfunction score as the mean integrated change in protein concentration, localization, and aggregation across genes assigned to each hallmark (normalized to young cells). Curves show progressive strengthening of hallmark-associated dysfunction with age.

Most of these hallmarks-of-aging-linked yeast proteins have human orthologs implicated in diverse diseases (Table S3), consistent with the view that these factors perform core cellular functions and that their age-associated destabilization may contribute to widespread dysfunction manifested as hallmarks of aging. Comparing these yeast proteins with human aging transcriptomes showed that 57.3% of their corresponding human orthologs show age-dependent mRNA abundance shifts (Table S3)^156–158^. Given the known decoupling between transcripts and protein abundance during aging, we additionally compared this set with a recently published multi-organ human aging proteomics dataset^159^. Among the human orthologs of these hallmark-linked yeast proteins, 91.6% showed age-associated abundance remodeling at the protein level (Figure 7B, Table S3). Enrichment analysis of these aging-sensitive human orthologs highlighted ribosome biogenesis/nucleolar pathways and mitochondrial functions (e.g., Fe-S biogenesis) (Figure 7C, S8I), supporting the idea that ribosome biogenesis and nucleolar function represent conserved, aging-sensitive nodes. Collectively, these cross-species comparisons are consistent with longstanding models proposing that core molecular features of aging are broadly conserved across evolution.

### Summary of molecular conduits linking hallmarks and their temporal emergence

Beyond established cross-hallmark links (e.g., NAD⁺ metabolism connecting mitochondrial function to chromatin regulation^1,2^), the atlas reveals additional molecular “conduits” that plausibly transmit dysfunction from one hallmark to another (Figure 7A, central inset). For example, loss of mitochondrial import/retention of shared base-excision repair factors (Ogg1, Ung1, Ntg1/2, Apn1, Cdc9) offers a direct route by which proteostasis and import defects can accelerate mtDNA instability. Similarly, reduction and aggregation of the deubiquitinase Ubp10, which deubiquitinates histone H2B and RNA polymerase I, provides a mechanism by which proteostasis collapse may propagate to epigenetic dysregulation and impaired ribosome biogenesis. In parallel, broad remodeling of nutrient transport and sensing programs predicts metabolic shift in mitochondria, while failure of mitochondrial Fe–S biogenesis is poised to compromise Fe–S–dependent nuclear genome maintenance^160–162^. In addition to these examples, the interconnectivity among the molecular functions of these hundreds of hallmark-linked proteins defines a systematic, proteome-scale map of inter-hallmark connectivity (Figure 7D, Table S4).

In parallel with the inter-hallmark connectivity map, we integrated age-associated shifts in protein concentration, localization, and aggregation to quantify molecular dysfunctions across hallmark-linked proteins and summarize these measures into hallmark-level temporal trajectories. Beyond early autophagy impairment, which is known to occur early due to loss of vacuolar acidification^5,106^, this analysis indicates that other hallmark programs also initiate early and progressively deepen, albeit at different rates, with nucleolar disruption, proteostasis decline, and mitochondrial dysfunction progressing more rapidly than other hallmark programs (Figure 7E). Notably, this meta-analysis resolved two hallmark groups: nucleolar disruption, proteostasis decline, and mitochondrial dysfunction exhibited similarly steep trajectories (Group 1), whereas epigenetic alterations, genomic instability, disabled autophagy, and deregulated nutrient sensing progressed more gradually and clustered together (Group 2). The separation between Group 1 and Group 2 trajectories was evident from the early age and persisted across the lifespan, suggesting that hallmark kinetics diverge early rather than only at late stages (Figure 7E). Because Group 1 hallmarks progress more rapidly, this pattern raises the possibility that early Group 1 dysfunction may contribute to the emergence or deepening of Group 2 hallmarks, potentially through their extensive functional interconnections (Figure 7D).

## Discussion

Replicative aging is often discussed through the framework of distinct hallmarks, yet how these hallmarks arise and connect with each other has remained unclear. We propose that progressive erosion of spatial proteome organization—loss of compartmental confinement, protein relocalization, shifts in local concentration, and aggregation—constitutes a unifying failure mode that disrupts spatially organized reactions, drives hallmark phenotypes, and spatiotemporally links these hallmarks. These age-associated changes are intrinsically spatial and highly heterogeneous, and thus are frequently obscured in bulk measurements. In this view, hallmark-linked dysfunction reflects not only altered expression programs but failures in where proteins reside, whether they remain soluble and available to engage substrates, and whether compartments can sustain specialized reaction environments.

An important implication from this multidimensional spatial proteome analysis is that molecular remodeling can be detectable well before overt failure in canonical functional assays. The temporal trajectories suggest that hallmark programs emerge early and deepen progressively, yet specific terminal functional collapse may lag because cells buffer stress through redundancy, feedback, and metabolic plasticity. For mitochondria, prior work based on membrane-potential collapse suggests apparent functional maintenance until mid-age^5^; whereas other studies suggest that a subset of aging cells enters alternative aging modes characterized by early mitochondrial heme-production dysfunction^23^. Our trajectory analysis indicates that mitochondrial deterioration begins earlier at the molecular level, with different mitochondrial pathways declining at distinct rates. Together, these observations are consistent with threshold-like behavior: graded molecular deterioration may accumulate for many divisions, while specific functional collapse becomes apparent only once buffering capacity is exceeded. Conceptually, this resembles the well-established “mitochondrial threshold effect” in heteroplasmy-driven disorders, where biochemical/clinical defects often emerge only once mutant mtDNA reaches a high fraction (commonly ∼60–80% or higher, depending on the mutation and tissue)^163^. Similar lags between early molecular remodeling and late functional endpoints may exist for other hallmarks, underscoring the value of multidimensional, spatially resolved measurements for identifying early vulnerabilities.

Within this framework, nucleolar dysfunction and ribosome-biogenesis remodeling emerge as a prominent early vulnerability. Rather than a uniform decline in ribosome-biogenesis factors, aging erodes nucleolar compartmentalization: processing, remodeling, and rRNA modification/surveillance components escape the nucleolus, become diffuse in the nucleoplasm, and frequently aggregate, while core rRNA transcription capacity persists longer. Ribosome biogenesis is one of the cell’s largest coordinated assembly programs and a dominant biosynthetic investment; accordingly, its failure is poised to make a major contribution to proteostasis decline. Moreover, because ribosomes produce essentially all cellular proteins, disruption of the spatial order of ribosome assembly could yield ribosomes with compromised translational capacity and fidelity, increasing proteotoxic load and amplifying demand on folding and degradation pathways—an instability that may push cells past a tipping point into persistent stress states. Once this node destabilizes, compensatory stress programs—translation repression, RNA rerouting, and remodeling of quality-control pathways—may transiently buffer proteostasis stress, but at the cost of accelerating downstream organelle dysfunction and metabolic drift. Building on prior observations that nucleolar size correlates with longevity across species and that translational fidelity strongly impacts lifespan^12,79–81^, together with the conserved age-associated remodeling of ribosome-biogenesis factors from yeast to human (Figure 7C), we propose that nucleolar dysfunction and ribosome-biogenesis defects merit consideration as an independent, conserved primary hallmark of aging.

The same spatial framework supports a network view of aging in which coupling can arise from inherent functional connectivity and shared limiting components—import capacity, proteostasis machinery, RNA-handling factors, and cofactors—across compartments that are highlighted by each hallmark. Shared factors can become selectively limiting within specific compartments—for example, when impaired mitochondrial import and proteostasis reduced the mitochondrial pool of shared DNA repair enzymes despite their nuclear abundance remained stable or even increased—providing a direct route from compartmental failure to mtDNA genome instability. This architecture provides a mechanistic rationale for why interventions that stabilize one hallmark often yield pleiotropic benefits: restoring spatial homeostasis at a key node can indirectly rebalance multiple downstream processes. Extending spatial proteomics across perturbations and species should enable principled maps of cross-hallmark causality and inform interventions that restore organization rather than targeting individual endpoints in isolation.

## Acknowledgments

We thank Dr. Maya Schuldiner and Dr. Michael Knop for providing SWAT libraries. We also thank the input and comments from lab members and colleagues, especially Catherine Chang. This work was supported by DP5OD024598, R21AG077556, Impetus grant, Larry L. Hillblom Foundation 2023-A-007-SUP, and R01AG075201 to C. Zhou and Hevolution Foundation to the Buck Institute for Research on Aging (HF-PART-23-1422047).

## Author contributions

C.Z. conceived and supervised the project. S.Y., L.L., and C.Z. designed the research. S.Y. and L.L. performed proteome-wide imaging data collection. L.V. analyzed protein localization, and L.L. analyzed proteome-wide protein abundance and concentration. C. Y., L.L., and L.V. performed data analysis and generated figures. S.Y. and M.W. contributed to phenotype identification. J.Z. and F.Z. generated plasmids and strains for localization validation. J.K.A provided support for some of the work. C.Z. drafted the manuscript with input from all authors.

## Declaration of interests

The authors declare no competing interest.

---

**Figure S1.**
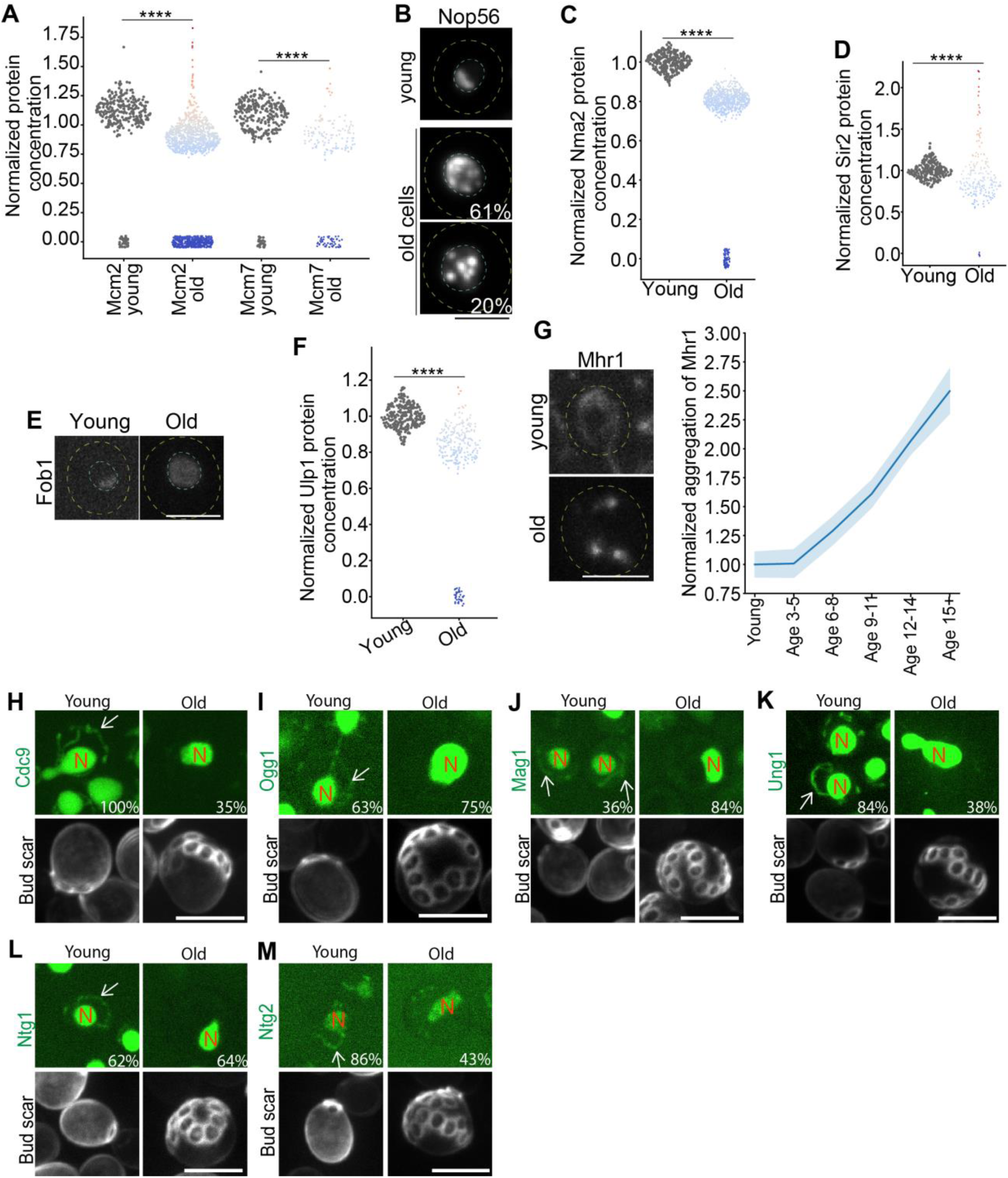
Additional data for age-associated molecular changes underlying genomic instability. (**A**) Quantification of protein concentrations for DNA replication-related proteins in young and old cells (≥15 generations; same definition used in other figures). Each dot represents an individual old cell, with values normalized to the mean value of the corresponding protein in young cells. Dot color indicates fold change relative to the young-cell mean (red: increase; blue: decrease; same in other figures) (**B**) Nucleolar morphological changes. Representative images of Nop56 localization and aggregation in young and old cells. The inset percentage indicates, for each age group, the fraction of cells exhibiting the phenotype shown in the representative images (same definition used in other figures). The cyan dashed line outlines the nucleus. (**C**) Quantification of protein concentrations for Nma2 in young and old cells. (**D**) Quantification of protein concentrations for Sir2 in young and old cells. (**E**) Representative images of Fob1 localization in young and old cells. The cyan dashed line outlines the nucleus. Fob1 redistributes from a crescent-shaped nucleolar pattern to a diffuse nucleoplasmic signal in old cells. (**F**) Quantification of protein concentrations for Ulp1 in young and old cells. (**G**) Representative images and quantification of Mhr1 aggregation during aging. Shown are mean and standard error (shade). Note that Mhr1 redistributes from mitochondria network into several puncta/aggregates. (**H-M**) Representative images and quantifications of localization for different DNA repair-related proteins in young and old cells. Arrow point to mitochondrial signal. N, nucleus. Scale bar: 5μm. See table S5 for number of cells quantified.

**Figure S2.**
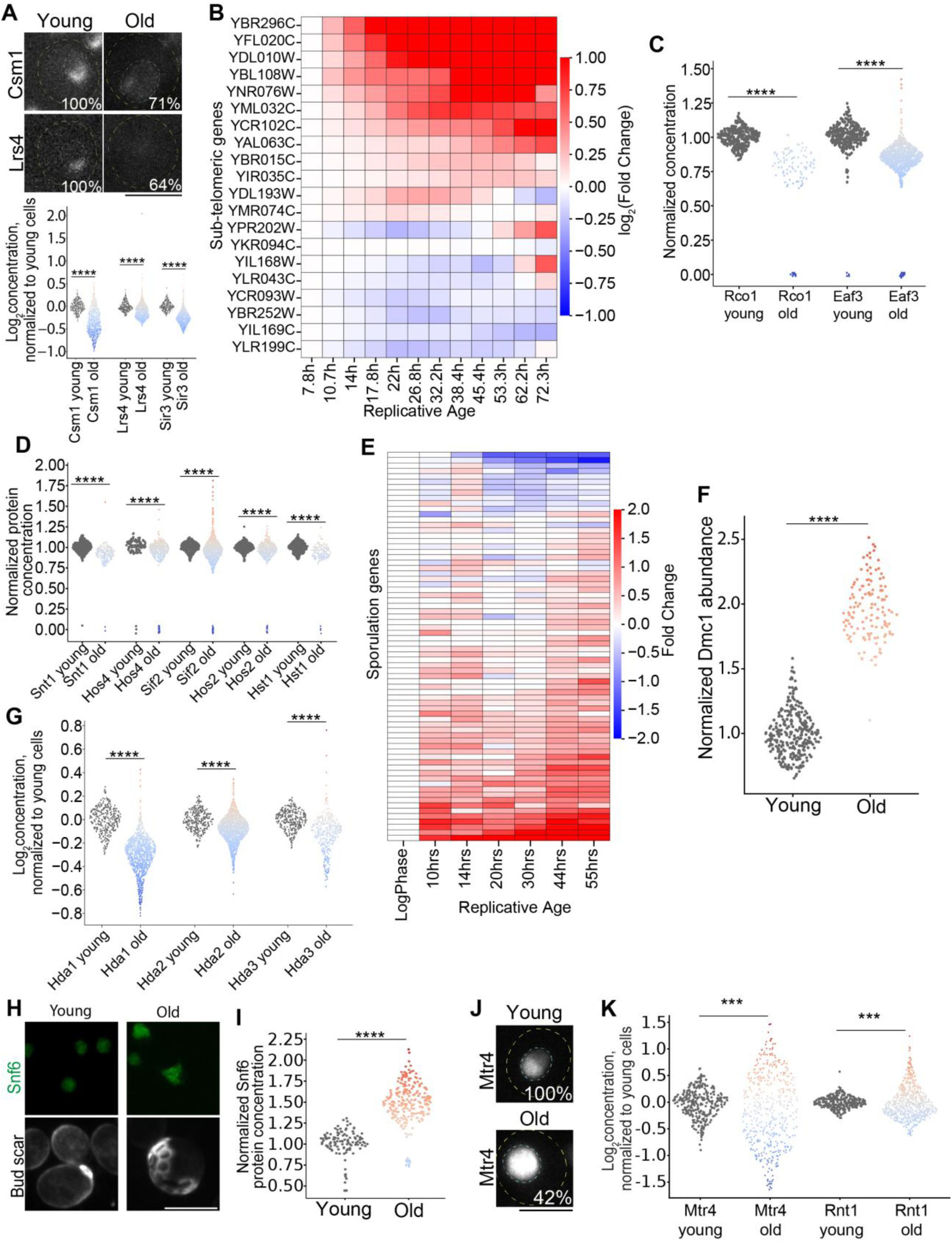
Additional data for age-associated molecular changes underlying epigenetic alterations. (**A**) Representative images and quantifications of protein concentrations for Csm1, Lrs4, and Sir3 in old cells (≥15 generations; same definition used in other figures). Each dot represents an individual old cell, with values normalized to the mean value of the corresponding protein in young cells. Dot color indicates fold change relative to the young-cell mean (red: increase; blue: decrease; same in other figures). The cyan dashed line outlines the nucleus (same for other figures). (**B**) Quantification of sub-telomeric gene expression by re-analyzing a published RNA-seq data^58^. (**C**) Quantification of protein concentration for subunits of Rpd3S complex in young and old cells. (**D**) Quantification of protein concentration for subunits of Set3C complex in young and old cells. (**E**) Quantification of sporulation gene expression by re-analyzing a published RNA-seq data^61^. (**F**) Quantification of protein abundance for Dmc1 in young and old cells. (**G**) Quantification of protein concentration for Hda1/2/3 in young and old cells. (**H-K**) Representative images and quantifications for Snf6, Mrt4, and Rnt1 in young and old cells. Scale bar: 5μm. See table S5 for number of cells quantified.

**Figure S3.**
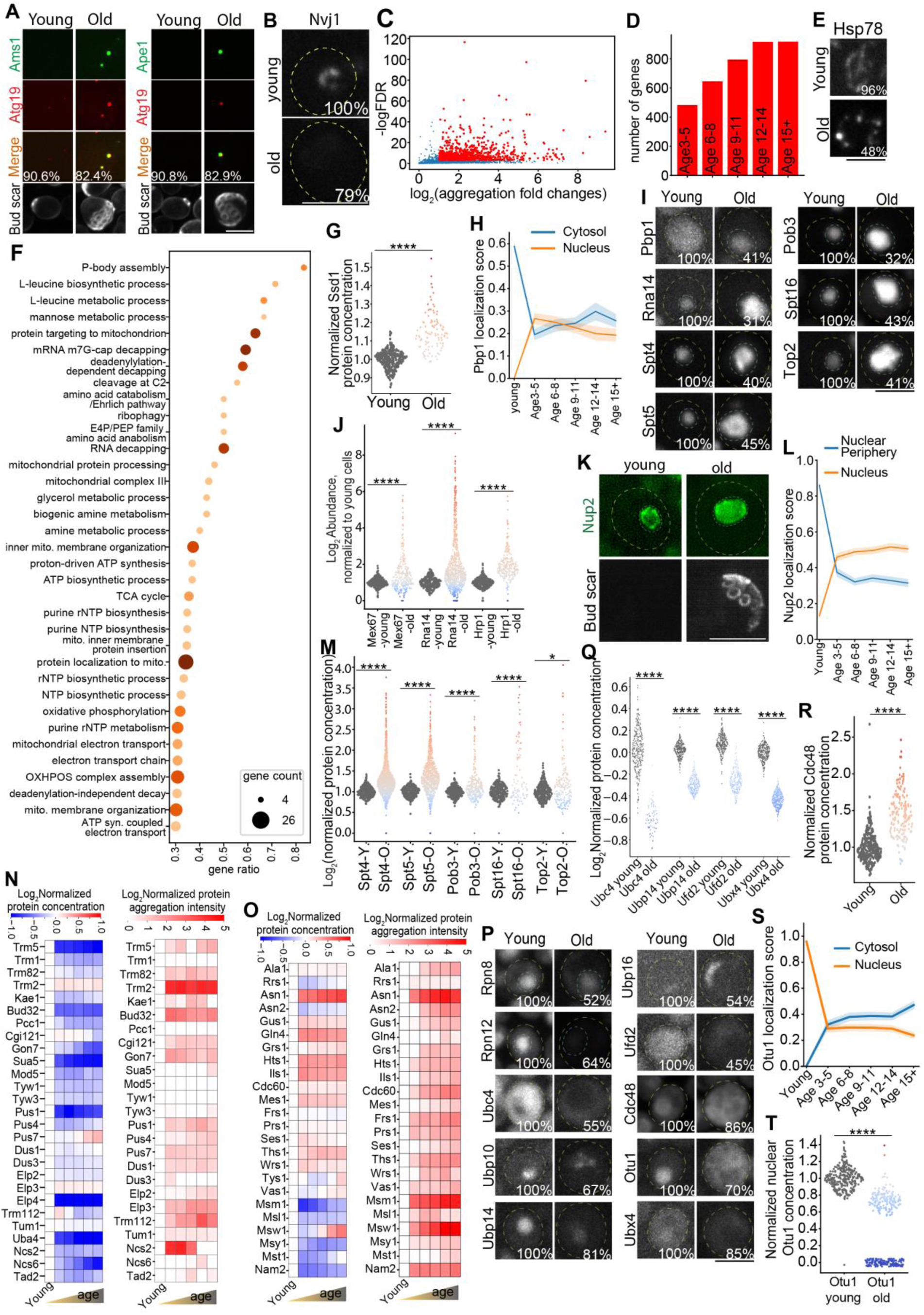
Additional data for age-associated molecular changes underlying disabled autophagy and loss of proteostasis. (**A**) Representative images showing the colocalization of Ams1 and Ape1 with Atg19 in the puncta formed in aged cells. (**B**) Representative images of Nvj1 localization and expression in young and old cells. The inset percentage indicates, for each age group, the fraction of cells exhibiting the phenotype shown in the representative images (same definition used in other figures). (**C**) Volcano plot summarizing proteome-wide quantification of age-associated increases in protein aggregation. Red points denote proteins showing >2-fold higher aggregation in cells aged ≥15 generations compared with the corresponding young cells. (**D**) Number of proteins exhibiting >2-fold increases in aggregation relative to the corresponding young cells across different aging stages. Note that many proteins started to aggregate as early as age 3-5. (**E**) Representative images and quantification of Hsp78 aggregation in young and old cells. (**F**) GO term biological process enrichment for the aggregating proteins during aging. Gene ratio is the number of genes from input list that fall into a give GO term divided by the GO term size. (**G**) Quantification of Ssd1 protein concentration in young and old cells (≥15 generations; same definition used in other panels). Each dot represents an individual old cell, with values normalized to the mean Ssd1 level in the corresponding young cells. Dot color indicates fold change relative to the young-cell mean (red: increase; blue: decrease; same in other figures). (**H**) Quantifications of Pbp1 localization changes during aging. Shown are mean and standard error (shade). (**I**) Representative images and quantification of localization changes of different proteins in young and old cells. (**J**) Quantification of Mex67, Rna14, and Hrp1 protein concentration in young and old cells. (**K-L**) Representative images and quantification of localization changes of Nup2 in young and old cells. Shown are mean and standard error (shade). (**M**) Quantification of Spt4, Spt5, Pob3, Spt16, and Top2 protein concentration in young and old cells. -Y., young; -O., old. (**N-O**) Quantification of protein concentration changes and aggregation during aging for tRNA modification enzymes (N) and tRNA aminoacyl synthetases (O). (**P**) Representative images and quantification of localization and expression changes of ubiquitin–proteasome pathway proteins in young and old cells. Notably, Ubp10 aggregated, Ubp16 relocalized from the cytosol to a discrete organelle-like structure, and Cdc48 and Otu1 shifted from the nucleus to the cytosol during aging. (**Q**) Quantification of protein concentration fold changes for Ubc4, Ubp14, Ufd2, and Ubx4 in old cells, normalized to the mean concentration in the corresponding young cells. (**R**) Quantification of Cdc48 protein concentration in young and old cells. (**S-T**) Quantification of localization (S) and expression (T) changes of Otu1 in young and old cells. Shown are mean and standard error (shade). Scale bar: 5μm. See table S5 for number of cells quantified.

**Figure S4.**
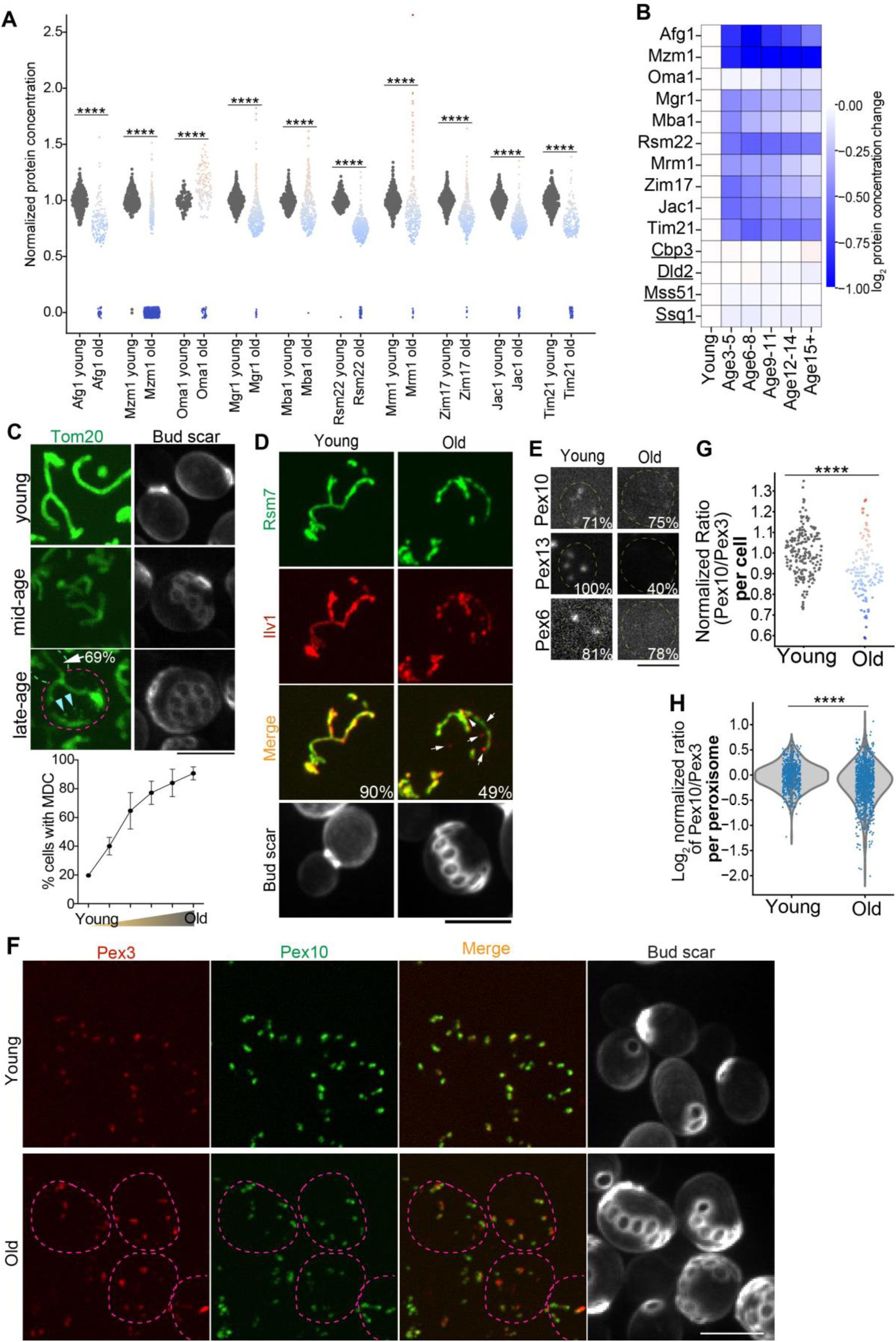
Additional data for loss of proteostasis within organelles. **(A)** Quantification of protein concentrations for different mitochondrial proteostasis machineries in young and old cells. Each dot represents an individual old cell, with values normalized to the mean protein level in the corresponding young cells. Dot color indicates fold change relative to the young-cell mean (red: increase; blue: decrease; same in other figures). **(B)** Quantification of concentrations for mitochondrial proteostasis factors that showed age-associated changes, alongside underlined mitochondrial proteins whose expression remained stable across aging. **(C)** Representative images and quantification showing the age-associated formation of mitochondria-derived compartments. Cyan arrowheads indicate mitochondria-derived compartments, and the white arrow marks a daughter cell (green dashed outline) produced by an old mother cell (pink dashed outline). Among old cells that undergo budding, 69% of their daughter cells did not inherit mitochondria-derived compartments. N= 348 old cells. **(D)** Representative images and quantification showing heterogeneous distribution of mitochondrial matrix proteins in aged cells. White arrows mark mitochondrial regions with distinct composition of the two matrix markers (red- or green-only signal). N=169 young cells and 148 old cells. **(E)** Representative images and quantification of different peroxisome proteins in young and old cells. **(F**, **G)** Representative images and quantification showing an age-associated decrease in the Pex10-to-Pex3 ratio in aging mother cells (pink dashed outline). In (G), each dot represents the mean Pex10:Pex3 ratio across peroxisomes within a single cell. **(H)** Peroxisome-level quantification of the Pex10-to-Pex3 ratio in young and aged cells. Each dot represents a single peroxisome. Note that aging increases inter-peroxisome variability in this key protein-import machinery. Scale bar: 5μm. See table S5 for number of cells quantified.

**Figure S5.**
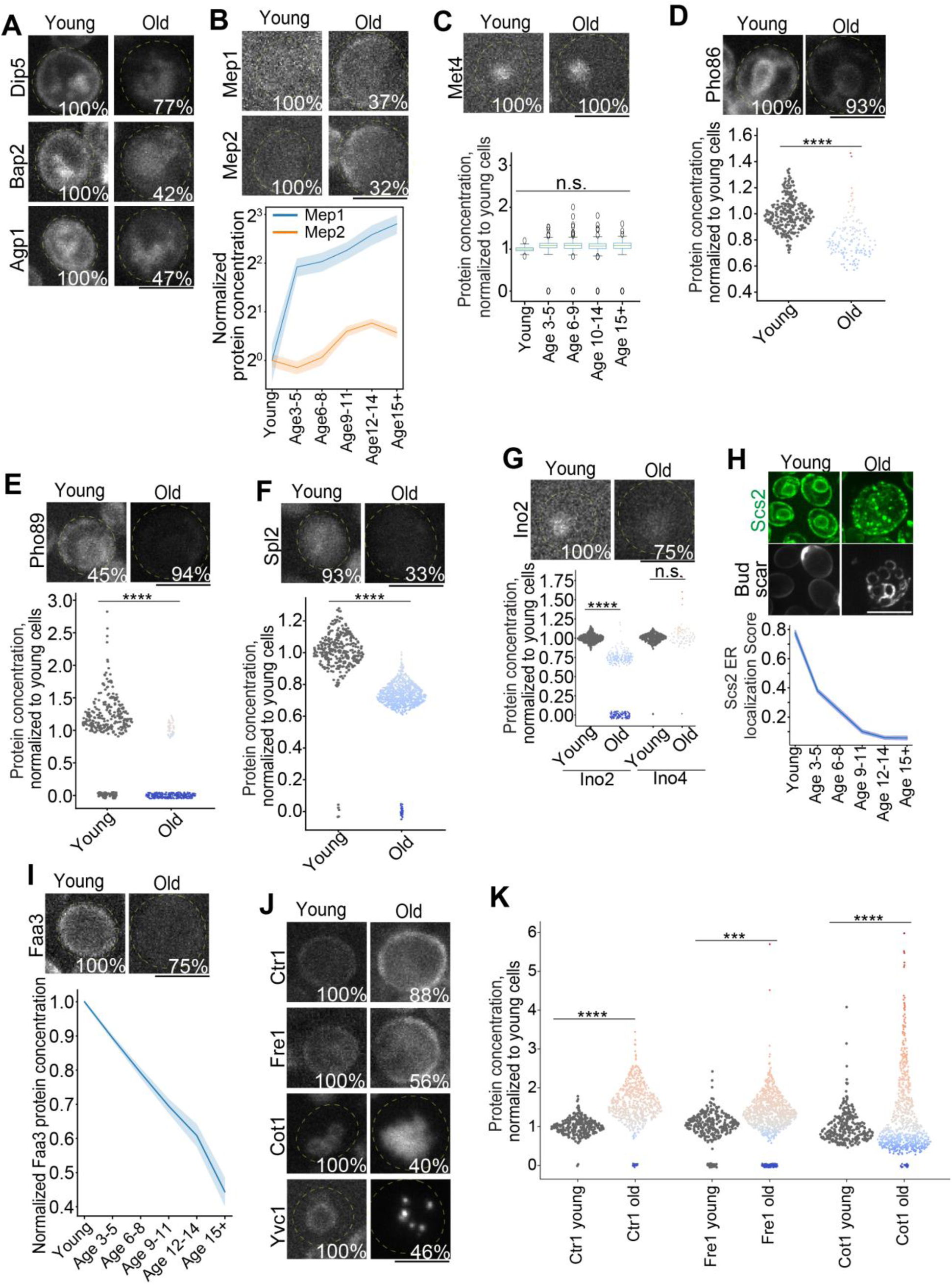
Additional data for age-associated molecular changes underlying deregulated nutrient sensing. **(A)** Representative images and quantification of protein concentrations for different amino acid transporters in young and old cells. The inset percentage indicates, for each age group, the fraction of cells exhibiting the phenotype shown in the representative images (same definition used in other figures). **(B)** Representative images and quantification of expression changes of Mep1/2 during aging. Shown are mean and standard error (shade). **(C)** Representative images and quantification of expression changes of Met4 in young and aged cells. **(D-G)** Representative images and quantification of protein concentrations for Pho86 (D), Pho89 (E), Spl2 (F), and Ino2/4 (G) in young and old cells. Each dot represents an individual old cell, with values normalized to the mean protein level in the corresponding young cells (same for other panels). Dot color indicates fold change relative to the young-cell mean (red: increase; blue: decrease; same in other figures). **(J)** Representative images and quantification of ER localization changes for Scs2 during aging. Shown are mean and standard error (shade). **(K)** Representative images and quantification of protein concentration of Faa3 during aging. Shown are mean and standard error (shade). **(J-K)** Representative images and quantification of different ion transporters in young and old cells. Scale bar: 5μm. See table S5 for number of cells quantified.

**Figure S6.**
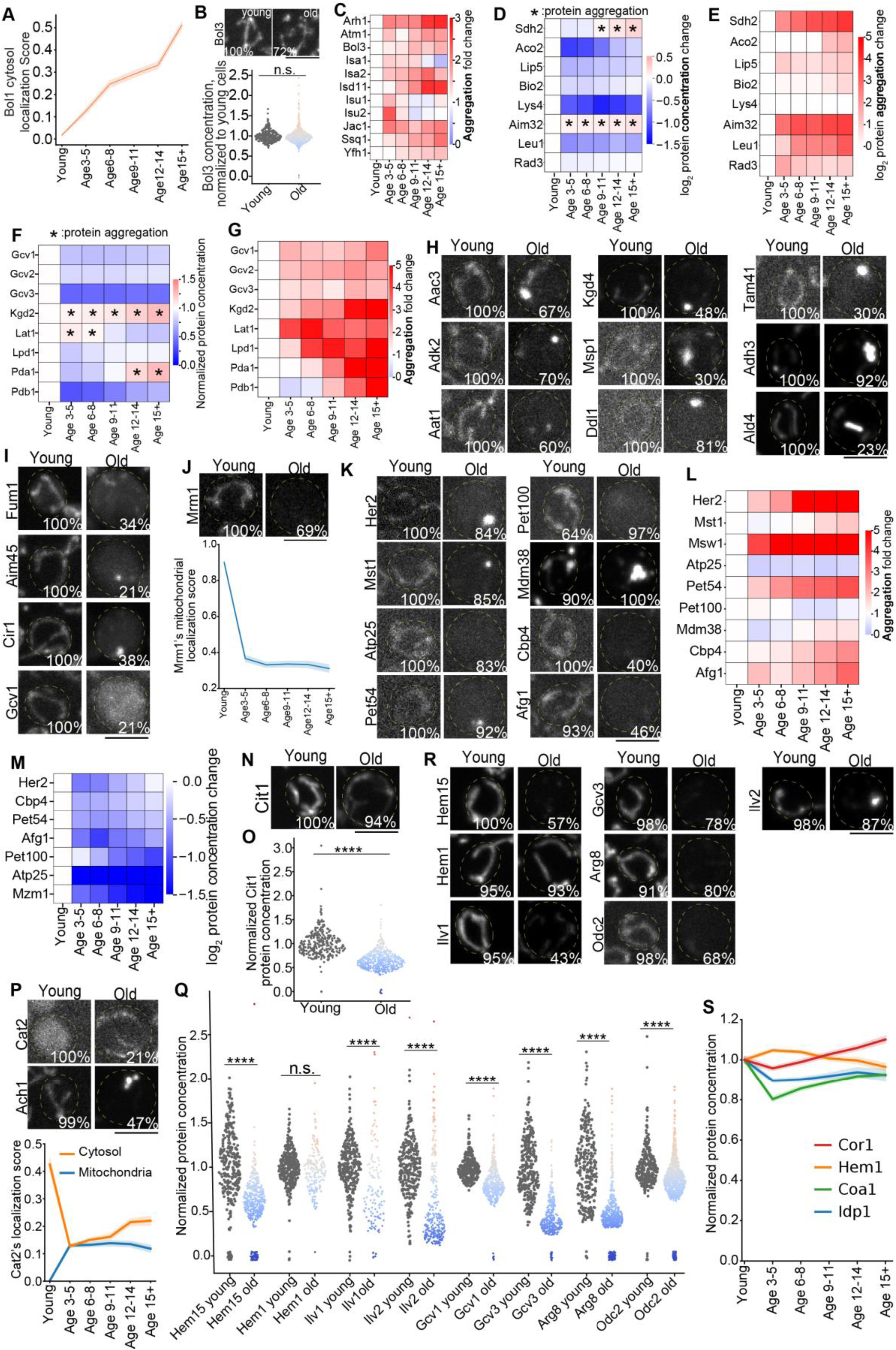
Additional data for age-associated molecular changes underlying dysfunctional mitochondria. **(A)** Quantification showing Bol1, a mitochondrial protein, relocalizes to cytosol during aging. Shown are mean and standard error (shade). **(B)** Representative images and quantification of protein concentrations for Bol3 in young and old cells. Each dot represents an individual old cell, with values normalized to the mean protein level in the corresponding young cells (same for other panels). Dot color indicates fold change relative to the young-cell mean (red: increase; blue: decrease; same in other figures). The inset percentage indicates, for each age group, the fraction of cells exhibiting the phenotype shown in the representative images (same definition used in other figures). **(C)** Quantification of protein aggregation for mitochondrial ISC pathway components across aging. **(D-E)** Quantification of protein concentration (D) and aggregation (E) for mitochondrial metabolic enzymes that depend on ISC-derived cofactors during aging. **(F-G)** Quantification of protein concentration (F) and aggregation (G) for mitochondrial metabolic enzymes that depend on lipoamide cofactor during aging. **(H)** Representative images and quantification of protein aggregation for different mitochondrial metabolic enzymes and structural proteins in young and old cells. **(I)** Representative images and quantification of protein localization changes for different mitochondrial proteins in young and old cells. **(J)** Representative images and quantification of Mrm1 in young and old cells. Shown are mean and standard error (shade). **(K-M)** Representative images (K) and quantification of protein aggregation (L) and concentration (M) for mitochondrial translation fidelity and assembly factors during aging. **(N-O)** Representative images and quantification of protein concentration for Cit1 in young and old cells. **(P)** Representative images and quantification of protein localization changes for Cat2 and Ach1 in young and old cells. Shown are mean and standard error (shade). **(Q-R)** Representative images and quantification of protein concentration for mitochondrial heme synthesis, amino acid catabolism, and acetyl-CoA metabolism in young and old cells. **(S)** Quantification of protein concentration for different mitochondrial proteins that remain stable during aging. Shown are mean and standard error (shade). Scale bar: 5μm. See table S5 for number of cells quantified.

**Figure S7.**
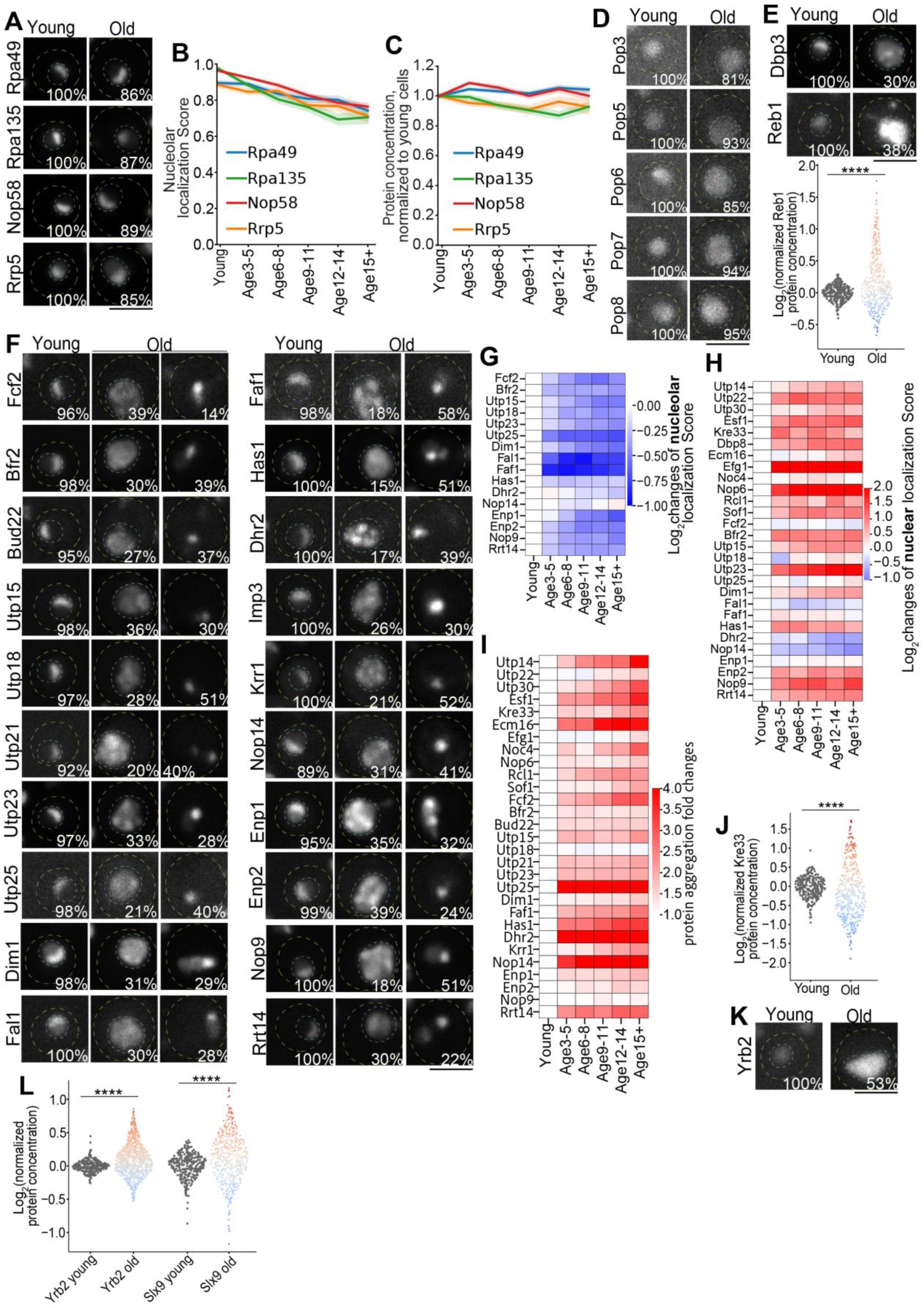
Additional data for age-associated changes in ribosomal small subunit (SSU) biogenesis. **(A**-**C)** Representative images (A) and quantification of nucleolar localization (B) and protein concentration (C) for SSU biogenesis factors that remain relatively stable during aging. The inset percentage in (A) indicates, for each age group, the fraction of cells exhibiting the phenotype shown in the representative images (same definition used in other figures). **(D)** Representative images and quantification of RNase MRP proteins in young and old cells. **(E)** Representative images and quantification of Dbp3 and Reb1 in young and old cells. Each dot represents an individual old cell, with values normalized to the mean protein level in the corresponding young cells (same for other panels). Dot color indicates fold change relative to the young-cell mean (red: increase; blue: decrease; same in other figures). **(F-G)** Representative images (F) and quantification of nucleolar localization (G) for additional factors involved in SSU maturation in young and old cells. For old cells, examples of both nucleolar escape and aggregation are shown in (F). **(H-I)** Quantification of nuclear localization (H) and protein aggregation (I) changes for factors involved in SSU maturation during aging. **(J)** Quantification of Kre33 protein concentrations in young and old cells. **(K-L)** Representative images and quantification of protein concentrations for Yrb2 and Slx9 in young and old cells. Scale bar: 5μm. See table S5 for number of cells quantified.

**Figure S8.**
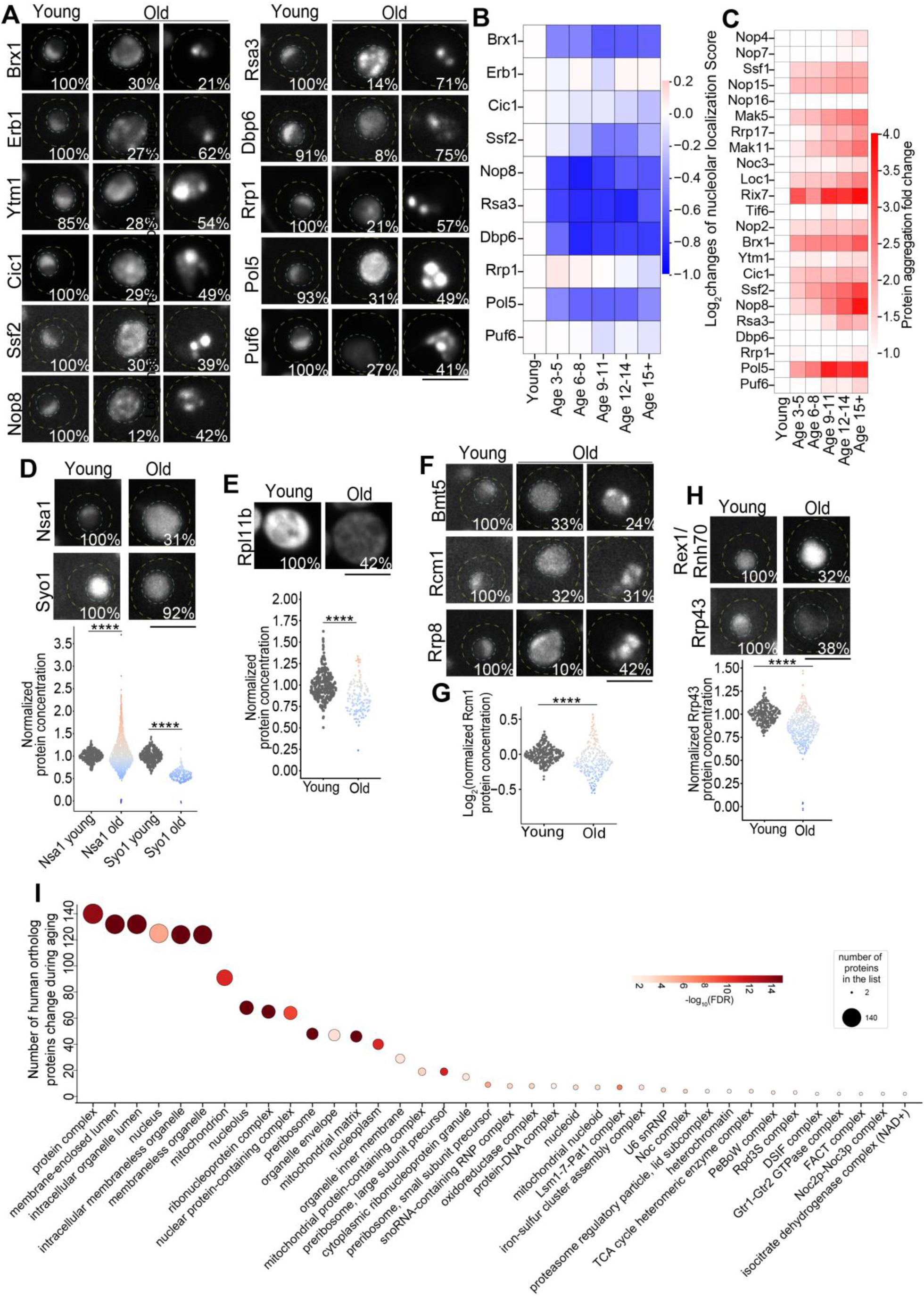
Additional data for age-associated changes in ribosomal large subunit (LSU) biogenesis. (A-B) Representative images (A) and quantification of nucleolar localization (B) for additional factors involved in LSU maturation in young and old cells. For old cells, examples of both nucleolar escape and aggregation are shown in (A). The inset percentage indicates, for each age group, the fraction of cells exhibiting the phenotype shown in the representative images (same definition used in other figures). **(C)** Quantification of protein aggregation changes for factors involved in LSU maturation during aging. **(D)** Representative images and quantification of protein concentrations for Nsa1 and Syo1 in young and old cells. Each dot represents an individual old cell, with values normalized to the mean protein level in the corresponding young cells (same for other panels). Dot color indicates fold change relative to the young-cell mean (red: increase; blue: decrease; same in other figures). **(E)** Representative images and quantification of Rpl11b protein concentrations in young and old cells. **(F-G)** Representative images of Bmt5, Rcm1, and Rrp8, and quantification of Rcm1 concentration in young and old cells. **(H)** Representative images and quantification of protein concentrations for Rex1/Rnh70 and Rrp43 in young and old cells. **(I)** Cellular component enrichment among the 219 human orthologs that show age-associated changes during human aging. Scale bar: 5μm. See table S5 for number of cells quantified.

## Yeast strains and culture condition

All *S. cerevisiae* strains, plasmids, primers, antibodies, and chemical reagents used in this study are listed in the Key Resources Table. Unless otherwise noted, strains are in the BY4741 genetic background. Genetic modifications were generated by PCR-mediated homologous recombination (Longtine et al., 1998)^167^ and verified by diagnostic PCR to confirm correct integration and the absence of aneuploidy at the chromosome carrying the targeted locus. mNeonGreen (mNG)-tagged strains were obtained from the seamless tagging collection (Meurer et al., 2018)^164^. Plasmids were constructed using Gibson Assembly (Gibson Assembly Master Mix; New England Biolabs, E2611S). For integration-based expression, linearized plasmids were integrated at the TRP1 locus.

Yeast cells were grown at room temperature in YPD (10 g/L yeast extract from BD, 20 g/L peptone from BD, and 20 g/L glucose), and refreshed for additional 2-3 hrs before imaging. The OD_600_ of cell culture was maintained between 0.4–1.0 overnight and throughout the experiments to avoid metabolic shifts caused by glucose exhaustion. All mediums used in the study were prepared by autoclaving the yeast extract and peptone for 20 min before adding the filtered carbon source as indicated.

## Confocal microscopy

Cells were imaged in 384-well glass-bottom plates (Cellvis, P384-1.5H-N, USA). Images were acquired on a Nikon CSU-W1 SoRa spinning-disk confocal microscope equipped with a 60×/1.27 NA Plan-Apochromat water-immersion objective (Plan Apo IR 60× WI DIC N2) and ORCA-Fusion BT cameras. Z-positioning was controlled by a piezo Z stage. Excitation at 405/488/561/561 nm was used for Calcofluor White, mNeonGreen (mNG), mScarlet3, and WGA, respectively, and emission was collected through appropriate filter sets (455/50 nm for Calcofluor White, 520/40 nm for mNG, 605/52 nm for mScarlet3, and 620/60 nm for WGA). SoRa mode with dual-camera acquisition was used to improve lateral resolution and reduce channel misalignment during co-localization analysis of dual-labeled strains. Image processing was performed in Fiji/ImageJ (NIH, Bethesda, MD). Cells from multiple repeats were quantified and shown.

## Automated Isolation of Cells with Advanced Replicative Age

Replicative old mother cells were enriched using a cell wall biotinylation and magnetic capture workflow executed on a Tecan Fluent 480 automated liquid-handling robot. Mid-log cultures (total input equivalent to ∼OD₆₀₀ 1.5) were dispensed into each well of 96-deep-well plates and pelleted by low-speed centrifugation. Cells were rinsed three times with PBS (pH 8.0), then resuspended in PBS (pH 8.0) and reacted with NHS–dPEG®12–biotin (0.35 mg per well; Sigma, QBD10198) for 30 min at room temperature with periodic mixing. Unreacted biotin reagent was removed by four washes with PBS (pH 7.2).

Biotinylated cells were subsequently incubated with streptavidin-coated magnetic beads (BioMag; 15 μL beads per strain; Polysciences, 84660-5) for 30 min at room temperature with intermittent mixing to allow binding. Bead-associated cells were retrieved on a 96-well magnetic stand and exchanged into fresh YPD medium. Replicative aging was then performed directly on the robot deck for 23 hours with automated medium refresh every 2 hours. To minimize settling and bead clumping during the aging period, cultures were gently resuspended by periodic air-bubble mixing (every 10 min). At the end of the aging run, bead-bound mother cells were recovered and washed three times to reduce carryover of newly budded daughters.

For imaging, cells were stained with CF®594-conjugated wheat germ agglutinin (WGA594) at 1 μg/mL for 1 hour to label bud scars, followed by Calcofluor White staining to delineate the cell wall (0.7 μL/mL of a 1 mg/mL stock; Sigma-Aldrich, 18909-100ML-F; 10 min incubation). For dual-labeled strains, only Calcofluor White staining was applied. With the exception of the initial low-speed centrifugation step prior to bead capture, all wash, labeling, staining, culture, and media-exchange operations were carried out using the Tecan Fluent 480 robot.

## Deep learning models for quantifying subcellular localization

Protein subcellular distribution was mapped at single-cell resolution using a dual-stream ensemble framework that integrates 2D projection-based features with 3D volumetric context (described in the accompanying study, Yoo et al., 2026a). The architecture comprises two parallel pathways: (i) a 2D CNN (DeepLoc) adapted to yeast morphology via transfer learning, and (ii) a 3D ResNet-50 backbone optimized for 8 × 64 × 64 voxel tensors. By combining maximum-intensity projections (MIPs) with axial-aware residual bottleneck blocks, the framework balances computational efficiency with the capacity to resolve complex spatial topologies. The classifier outputs 17 categories, including 16 organelle localizations and a “None” class used for quality control to flag unexpressed cells.

Model outputs from the two streams were integrated to quantify localization. For each cell, raw logits were transformed into class probabilities using a softmax function, and a consensus probability vector was computed as the unweighted arithmetic mean of the two streams. The resulting 17-class probability signature enables high-confidence assignment of subcellular residency based on the maximum consensus probability. The classifier was applied to 79.4 million cells across ∼5,661 strains. Cells with “None” class probability > 0.6 were excluded to remove low-expression or non-expressing cells.

To quantify age-dependent proteome redistribution across the replicative lifespan, quality-filtered cells were partitioned into six age bins (Young, 3–5, 6–8, 9–11, 12–14, and 15+), defined by WGA-stained bud-scar counts. Single-cell measurements were summarized as strain-specific “morphological centroids,” defined as the mean 17-dimensional probability vector for each age bin. Age-bin cohorts with fewer than 50 cells were excluded to ensure statistical power.

## Quantification of protein concentration

For each single-cell mNeonGreen (mNG) crop, voxels outside the corresponding cell mask were blanked (set to zero) throughout the z-stack. Within the masked region, organelle-enriched signal was separated from diffuse cytosolic signal using two alternative segmentation strategies applied to the full 3D volume: (i) Yen intensity thresholding, and (ii) a percentile rule in which the brightest 6% of voxels were designated as organelle signal. For each strategy, the mean intensity ratio between the putative organelle compartment and the remaining cytosolic compartment was computed, and the strategy yielding the larger ratio was selected on a per-cell basis.

Compartment assignments from our ensemble localization classifier were used to standardize downstream quantification. Cells classified as cytoplasm were treated as lacking a distinct organelle compartment for this analysis: any segmented organelle mask/intensity was reassigned to the cytosolic compartment (i.e., organelle mask and intensity were set to zero). Cells classified as none were excluded by setting both organelle and cytosolic masks/intensities to zero.

As a quality-control step, the fraction of the cell occupied by the organelle mask was calculated. When the organelle mask exceeded 20% of the cell area/volume—most commonly due to slight misregistration between the mNG stack and the cell mask that introduced background and distorted thresholding—the original mNG volume was re-segmented using a two-step procedure: a coarse top-50% intensity cutoff to remove dark background, followed by the top-6% cutoff to define organelle-enriched signal. Cells classified as vacuole or vacuolar membrane were exempt from this re-thresholding because these compartments can legitimately occupy >20% of the cell.

For quantification, protein abundance was defined as the sum of mNG signal within the final organelle mask, whereas protein concentration was defined as the mean mNG intensity within the organelle mask.

## Quantification of protein aggregation

As aggregate detectability varies across organelle patterns, three complementary approaches were implemented for puncta-based aggregate detection. For consistency, all aggregate quantification was performed on z-maximum–projected mNG images. Dot-like organelles (Golgi, lipid particles, endosomes, spindle pole, bud neck, peroxisomes, and actin) were excluded because their native punctate morphology confounds aggregate calling.

Method 1 (high-sensitivity patterns). Applied to strains with cell-periphery signal and ER patterns with mean mNG intensity >560. mNG stacks were z-max projected, and pixels outside the cell mask were set to zero. Yen thresholding was applied to the projected image to define organelle-associated signal. Pixels with intensity >2× the mean organelle intensity were segmented as puncta, and objects <15 pixels were removed as noise.

Method 2 (medium-sensitivity patterns). Applied to mitochondrial patterns and ER patterns with mean mNG intensity of 280–560. Puncta were detected on z-max projections using a Laplacian-of-Gaussian (LoG) detector (min_sigma = (1.2, 1.5), max_sigma = (2, 3), num_sigma = 30, threshold = 0.03). Neighboring puncta were merged if they overlapped by >10% of pixels. To reduce false positives, only puncta with mean intensity in the top 5% among all cells for that strain were retained as aggregates.

Method 3 (low-sensitivity patterns). Applied to nucleolus, nucleus, cytoplasm, vacuole, vacuolar membrane, nuclear periphery, and ER patterns with mean mNG intensity <280. Puncta were detected on z-max projections using LoG (min_sigma = (1.2, 1.5), max_sigma = (2, 2.4), num_sigma = 30, threshold = 0.03), then filtered by (i) radius >1.7 and (ii) intensity enrichment relative to local background, defined as mean_intensity / mean_intensity of a surrounding annulus (radius = 2.5× punctum radius) >1.65.

For all methods, aggregate abundance was quantified as the total mNG signal within the segmented puncta regions

## Localization map

Global dynamics of proteome remodeling were visualized by projecting 17-dimensional strain centroids into a two-dimensional landscape using t-distributed stochastic neighbor embedding (t-SNE). Embedding was performed with scikit-learn using a cosine distance metric, which emphasizes similarity in the direction of localization signatures rather than absolute magnitude. A perplexity of 30 was used to preserve local neighborhood structure while providing an interpretable view of the global organization of the dataset.

To enable standardized temporal comparisons, all age-stratified cohorts were embedded within a shared global coordinate system, allowing age-dependent shifts to be assessed without artifacts introduced by independent embeddings. Within this landscape, territories corresponding to each organelle class were delineated using convex hulls, providing fixed spatial benchmarks for the 16 subcellular compartments.

Age-dependent trajectories were visualized by tracking the movement of strain centroids relative to these compartment boundaries across successive age bins. This trajectory-based representation captures both discrete translocation events and more gradual shifts in spatial organization, enabling high-resolution analysis of how proteome localization landscapes change over the course of replicative aging.

## Quantitative assessment of organelle rejuvenation

To quantify structural recovery across organelles, a rejuvenation score was calculated to represent the fraction of the age-associated localization shift that is avoided (or reversed) in age-0 cells. For each of 16 organelle categories, the mean L1 (Manhattan) distance from the young baseline (sum of absolute differences across features) was computed for both age-0 cells and age-15+ old mother cells. The rejuvenation score was then defined as:

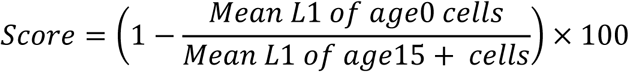

In this framework, a score of 100% signifies a complete return to the young cell localization baseline, while a score of 0% indicates a structural state indistinguishable from the age-15+ old mother cells. Variability was assessed using the Standard Error of the Mean (SEM), propagated to the final score to reflect the precision of the estimated recovery.

## Quantification of peroxisome protein heterogeneity

Peroxisome proteins of interest (Pex10, Pex13, Pex19) were endogenously tagged with mNeonGreen, and each was co-expressed with Pex3-mScarlet (a peroxisome marker that remains stable during aging). Both mNeonGreen and mScarlet z-stacks were max-intensity Z-projected. mScarlet max-intensity projections were thresholded using the Yen algorithm to generate a binary mask, followed by watershed segmentation to separate individual objects (peroxisomes). For each segmented peroxisome, mean mNeonGreen and mScarlet intensities were measured. The mNeonGreen/mScarlet ratio was then calculated per peroxisome (summarized at the single-cell level) and compared across ages to quantify peroxisome-to-peroxisome heterogeneity.

## Inter-hallmark connectivity map

A genome-wide functional interaction network was downloaded from GeneMANIA (Saccharomyces cerevisiae combined network; file: COMBINED.DEFAULT_NETWORKS.BP_COMBINING.txt)^168^. Pairwise gene–gene interaction weights were extracted for all hallmark-associated genes (Table S4). For visualization, only the top 0.1% strongest inter-hallmark connections (i.e., gene pairs assigned to different hallmarks) were plotted. Hallmark–hallmark connectivity was quantified as the sum of interaction weights across all inter-hallmark gene pairs between two hallmark gene sets.

We define hallmark–hallmark connectivity between hallmark gene sets A and B as:

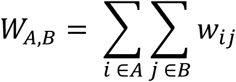

where *W_ij_* is the GeneMANIA pairwise interaction weight between genes i and j.

## Hallmark-level temporal trajectories during replicative aging

Genes associated with each hallmark of aging were curated (Table S4). For each gene and age group, we computed an integrated “dysfunction” score by combining changes in subcellular localization, protein concentration, and aggregation relative to young cells.

### Localization change

Localization remodeling was quantified as the sum of absolute differences across the 17 compartment localization scores between young and the indicated age group:

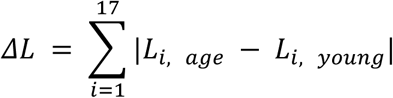

### Protein concentration change

Protein concentration was quantified as the fold change (FC) in organelle mean intensity between the age group and young (age/young). To treat increases and decreases symmetrically, fold changes < 1 were converted to their reciprocal:

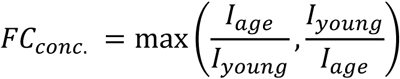

### Aggregation change

Aggregation was quantified as the fold change in aggregate abundance between the age group and young:

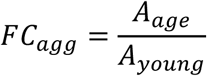

Integrated dysfunctional score. The per-gene integrated score was computed as:

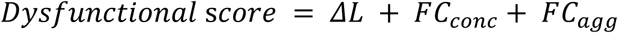

For each hallmark and age group, we computed the hallmark-level dysfunctional score as the mean integrated score across all genes assigned to that hallmark and normalized to the young cells before plotting these values across age to generate hallmark trajectories.

## Quantification and statistical analysis

Experiments were repeated multiple times to confirm reproducibility. In each experiment, strains were labeled by numbers. Image quantifications were done by batch processing without knowing the strain details. All quantifications are presented as the means ± standard error of mean (SEM). Statistical test using Mann-Whitney U test was included in each figure and figure legend. *: p<0.05; **: p<0.01; ***: p<0.001; ****: p<0.0001.

## Notes

### Competing Interest Statement

The authors have declared no competing interest.

